# Intragenomic conflicts with plasmids and chromosomal mobile genetic elements drive the evolution of natural transformation within species

**DOI:** 10.1101/2023.11.06.565790

**Authors:** Fanny Mazzamurro, Jason Baby Chirakadavil, Isabelle Durieux, Ludovic Poiré, Julie Plantade, Christophe Ginevra, Sophie Jarraud, Gottfried Wilharm, Xavier Charpentier, Eduardo P. C. Rocha

## Abstract

Natural transformation is the only mechanism of genetic exchange controlled by the recipient bacteria. We quantified its rates in 1282 strains of the human pathogens *Legionella pneumophila* (Lp) and *Acinetobacter baumannii* (Ab) and found that transformation rates evolve by large quick changes as a jump process across six orders of magnitude. Close to half of the strains are non-transformable in standard conditions. Transitions to non-transformability were frequent and recent, suggesting that they are deleterious and subsequently purged by natural selection. Accordingly, we find that transformation decreases genetic linkage in both species, which often accelerates adaptation. Intragenomic conflicts with chromosomal mobile genetic elements (MGEs) and plasmids could explain these transitions and a GWAS confirmed systematic negative associations between transformation and MGEs: plasmids and other conjugative elements in Lp, prophages in Ab, and transposable elements in both. In accordance with the modulation of transformation rates by genetic conflicts, transformable strains have fewer MGEs. Defense systems against the latter are associated with lower transformation except the adaptive CRISPR-Cas systems which show the inverse trend. The two species have different lifestyles and gene repertoires, but they exhibit very similar trends in terms of variation of transformation rates and its determinants, suggesting that genetic conflicts could drive the evolution of natural transformation in many bacteria.

## Introduction

Natural transformation consists in the uptake of exogenous DNA from the external surroundings of bacteria and its integration in their chromosome by homologous recombination. Unlike conjugation and transduction, two other key mechanisms of horizontal gene transfer (HGT), natural transformation is not encoded by mobile genetic elements (MGEs). It is the only known mechanism of HGT encoded and under the direct control of the recipient bacteria. It’s the mechanism behind the “transforming principle” that resulted in the discovery of HGT [1] and ultimately to the identification of DNA as the material of genetics [2]. Transformation requires a transient physiological state, the “competence” state, during which the machinery necessary for DNA import and integration in the chromosome is expressed [3]. Relatively few species have been demonstrated to be naturally transformable, but many more encode the necessary machinery and it is suspected that they are also transformable [4]. The process starts by the capture of exogenous DNA by an extracellular type IV pilus (Pil genes), whose retraction conveys the DNA at the cell surface [5]. In *Helicobacter* this step depends on a machinery derived from type 4 secretion system [6]. A nuclease converts the DNA into single-stranded DNA (ssDNA) before its transport to the cytoplasm by ComEC [3]. Once in the cytoplasm, the incoming ssDNA is protected from degradation by DprA, which also recruits RecA [3]. The recipient homologous recombination machinery is then involved in genetic exchanges with the bacterial chromosome. In diderms, the ComM protein has a key role in this process, facilitating genetic exchanges between long heterologous DNA sequences [7].

Even if competence for natural transformation was discovered almost a century ago, the reasons for its existence are still debated [8] (Figure S1). They include the promotion of allelic recombination [8], the acquisition of nutrients [9], and the uptake of DNA for repair [4]. The impact of transformation is significant. For instance, transformation enabled the acquisition of antibiotic resistance determinants by *Campylobacter jejuni* [10], *Streptococcus pneumoniae* [11] and *Acinetobacter baumannii* [12]. Recently, it was proposed that transformation eliminates deleterious MGEs by recombination in the flanking chromosomal core genes [13]. This *chromosome-curing* hypothesis implicates the existence of intragenomic conflicts between MGEs and the host regarding natural transformation. Accordingly, MGEs of *V. cholerae* (an integrative conjugative element, ICE) and of *C. jejuni* (a prophage) encode DNases that prevent transformation [14], [15]. Many other MGEs insert and disrupt key competence genes [13]. It is possible that several of these hypotheses contribute to selection for natural transformation.

The core components of the DNA uptake system and of homologous recombination are widely conserved across Bacteria. Yet, large variations of transformation frequencies have been observed and many species are described as non-transformable even though they encode all necessary components. For instance, *Pseudomonas stutzeri* is transformable whereas *Pseudomonas aeruginosa* is widely viewed as non-transformable [16]. Differences in transformability within species have also been observed. In well-established transformable species such as *S. pneumoniae*, *P. stutzeri* and *Haemophilus influenzae*, from 30% to 60% of the isolates [17]–[19] consistently failed to transform. Extensive variations were also reported in wildlife, clinical and human isolates of *Acinetobacter baumannii* [20] and in clinical isolates of *Legionella pneumophila* [21]. The reasons of such large within-species variations in transformation frequencies remain poorly understood. Of note, hypotheses explaining the existence of transformation do not necessarily explain large variations of their rates within species. For example, one does not expect huge variations in transformation between strains under similar growth conditions when there is selection for DNA repair or use of DNA as a nutrient. This does not imply that such hypotheses are incorrect, yet suggests that additional forces are at play. Notably, if intra-genomic conflicts affect transformation rates then low transformation rates could evolve even if they are deleterious to the bacterium [21].

The previously cited studies used relatively small samples (fewer than 150 and sometimes a few dozens), which precludes the use of powerful statistical methods to understand the genetic basis of phenotypic variation. Here, we characterize the transformation rates and identify their genetic determinants in two phylogenetically distant *Gammaproteobacteria* species, *L. pneumophila* (Lp) and *A. baumannii* (Ab). These are important pathogens with different characteristics. Ab is one of the most worrisome antibiotic resistant nosocomial pathogens [22]. Some strains are now resistant to nearly all antibiotics [23], an evolutionary process driven by chromosomal recombination events (possibly resulting from transformation) and by MGEs carrying antibiotic resistance genes [24]. Lp is an intracellular pathogen responsible for community-acquired severe pneumonia [25], where recombination and HGT drive the emergence of epidemic clones [26], but is not usually antibiotic resistant. Ab and Lp are therefore complementary models for revealing commonalities, and specificities, in the evolution of natural transformation. Here, we obtained two very large sets of genomes and transformation rates to characterize the distribution and evolutionary pace of these rates and to test if the trait is under selection. We use them to assess the hypothesis that genetic conflicts caused by MGEs contribute to explain variations in transformation rates.

## Results

### Variable transformation rates across Acinetobacter and Legionella *strains*

We analysed 496 draft genomes of Ab and 786 of Lp. The species pangenomes included 31,103 (Ab) and 11,932 (Lp) gene families. Ab genomes had on average 3598 genes with a coefficient of variation of 5.0%, whereas genomes of Lp had on average 3091 genes with a coefficient of variation of 3.3%. We used the gene families present in more than 95% of the genomes (persistent gene families: 2325 in Lp and 2629 in Ab) to build two types of recombination-aware phylogenetic trees for each species by maximum likelihood (see Methods, Table S1). The phylogenetic reconstructions and most analyses in this study were done on three pairs of phylogenetic trees (ignoring recombination, using random positioning of genes, and using the most likely organisation). Even if we identified many recombination tracts (see below), the qualitative results of all major analyses were similar in all the cases. Hence, only the recombination-aware method using the consensus genome organisation is presented in the text. The phylogenetic trees of the two species have approximately similar average root-to-tip distances (Ab: 0.031; Lp: 0.047 subst^-1^), but the Lp tree has many more small terminal branches than Ab. Altogether, the Ab dataset is more variable in terms of gene repertoires, but the species are of comparable age (distance to the species’ last common ancestor).

We used a luminescence-based assay to quantify the ability of each strain to acquire DNA by natural transformation (see Methods). Bacteria are provided with a linear transforming DNA consisting of the *Nluc* gene flanked by sequences homologous to the chromosome. Integration by natural transformation results in Nluc expression which is detected following addition of furimazine. Transformation assays were performed multiple times and revealed good concordance. Strains known to be non-transformable (Ab Δ*comEC*, Lp Lens) were used to define the minimal detection limit of the method, which is of the same magnitude in the two species: 300 for Lp and 400 for Ab. The distributions of the average values of transformation extend below the detection limit of transformation and have a long tail of higher values spanning several orders of magnitude (Figure 1B). As a result, 52% of the strains were deemed transformable in Lp and 64% in Ab. The range of transformation rates is higher in Ab (6 orders of magnitude) than in Lp (4).

**Figure 1.**
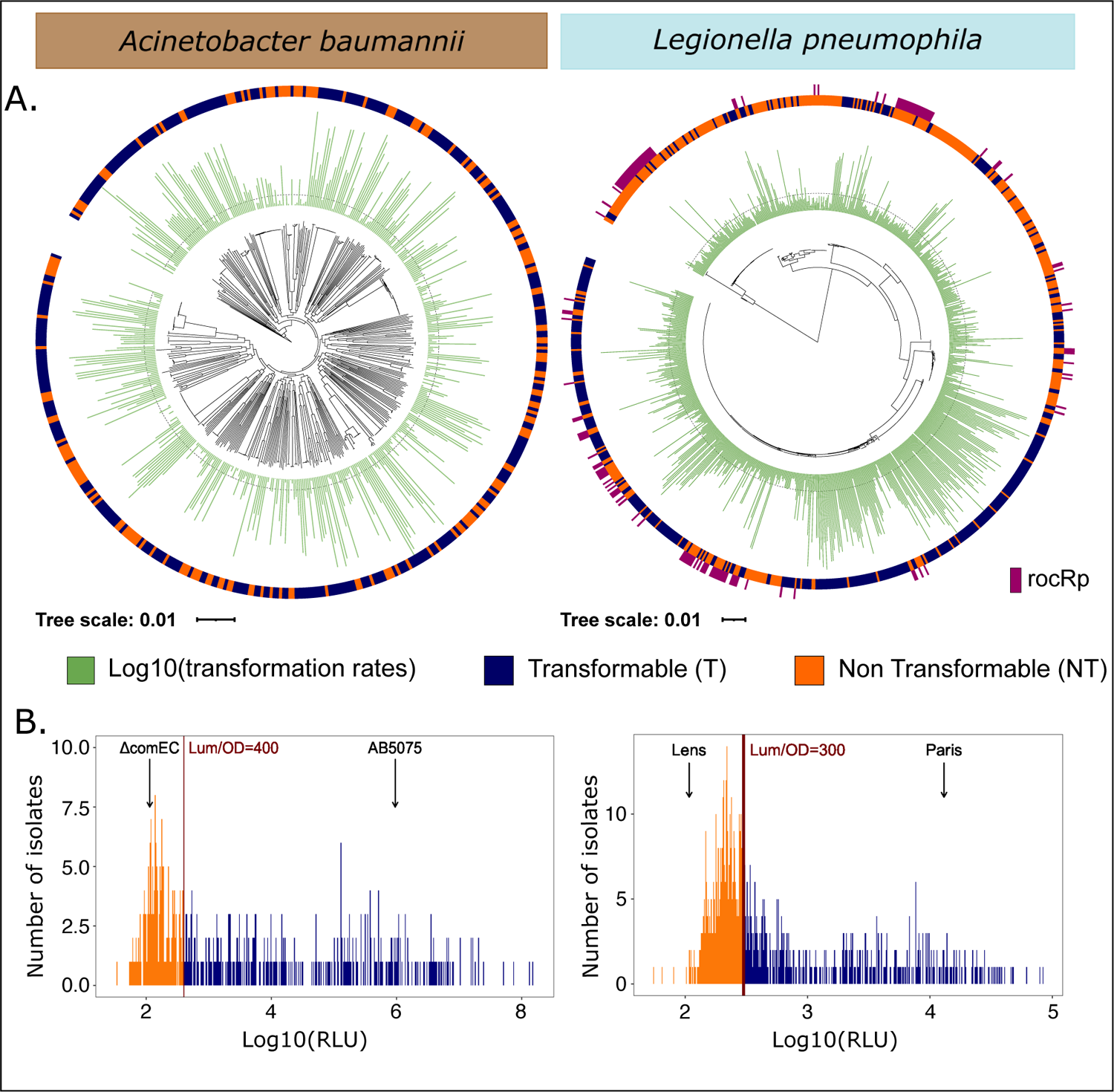
Distribution of the transformation phenotype. A. Distribution of the log10-transformed average transformation rates and of the binary trait across the recombination-free rooted phylogenetic trees of *Acinetobacter baumannii* (left) and *Legionella pneumophila* (right). sncRNA rocRp presence (dark pink) distribution is also represented across *Legionella pneumophila* phylogenetic tree. The dashed line in the log10-transformed average transformation rates distribution corresponds to the transformation rate threshold that separates transformable from non-transformable strains. B. Distribution of the log10-transformed average transformation rates in *Acinetobacter baumannii* (left) and *Legionella pneumophila* (right). The red vertical line stands for the threshold between transformable and non-transformable strains. ΔcomEC Ab strain and Lp Lens strain are known non-transformable strains. AB5075 Ab strain and Lp Paris strain are among the most transformable known strains.

Variations in transformation rates could be caused by differences between the focal strains and the template sequence used to produce linear transforming DNA (reference Ab A118 and Lp Paris strains). Instead, we found that transformation rates are highly variable across the phylogenetic trees (Figure 1A). The variations in transformation rates did not correlate with the strains’ phylogenetic distance (Ab: Spearman correlation ρ=0.0034, p=0.22; Lp: ρ=-0.046, p<2.2×10^-16^) even between closely related strains at a patristic distance of less than 0.02 nucleotide substitutions per site (Ab: ρ=0.029, p=0.18; Lp: ρ=-0.034, p<2.2×10^-16^) (Figure S2). The phylogenetic distance between the focal and reference strains are not correlated to the differences in transformation rates for the most distant strains (patristic distances larger than 0.04 nucleotide substitutions per site) (Ab: ρ=-0.011, p=0.81; Lp: ρ=0.029, p=0.80, Figure S3). Hence, transformation rates do not seem to depend on the phylogenetic distance between the donor and the recipient bacteria. Instead, they exhibit high variability across the species.

### Transformation rates evolve according to a jump process

We used the classical Pagel’s λ and Blomberg’s K indices to model the change of transformation rates through time, but they provided inconsistent results (Table S2). The Fritz and Purvis’s D statistic indicated that the binary trait is less clumped than expected under a Brownian model but more than if it was random (Lp, D=0.52 and in Ab D=0.58). This led us to assess how nine models with diverse evolutionary dynamics [27] fitted the evolution of the transformation rates (log transformed, Table S3). Six of them use Lévy processes with two components: a Brownian motion and a pure jump process.

The latter are stochastic processes making abrupt transitions between states where they remain for a given holding time. The best fitting models in both species were all jump processes (Jump Node, Variance Gamma, Brownian Model + Normal Inverse Gaussian, and Brownian Model + Variance Gamma, Table S3), which had all approximately similar fit with the data (similar AIC_c_) because they only differ in the type of random distribution defining the frequency and the amplitude of the jump. The good fit of these models suggests that natural transformation evolves by sudden changes separating periods of relative stability.

### Loss of transformation is counter-selected

The previous results suggest sudden transition between transformability and non-transformability. If transitions result from intragenomic conflicts, the latter must result from the deleterious impact of the loss of transformation for bacterial fitness. To test this hypothesis, we inferred the ancestral states of transformation in the phylogenetic tree and searched for recent transitions, i.e. those occurring in terminal branches. More than a fifth of the terminal branches had a phenotype transition (Lp: 20%; Ab: 20%), and there was a very large excess of transitions to non-transformability relative to those towards transformability (Lp: 79% of all events, *X*^2^, p<2.2×10^-16^; Ab: 81%, *X*^2^, p<2.2×10^-16^). If the process was at equilibrium there should be an equal number of transitions in both directions. Excess of transitions towards one of the states is usually a sign of purifying selection, i.e., a process where genetic changes enrich populations in a state (here, non-transformability) that is deleterious and therefore gradually purged by natural selection [28].

If the loss of transformation is deleterious for bacteria, then it should impact the evolutionary trajectories of bacterial lineages. Notably, if lineages of non-transformable strains are less fit then they are expected to be shorter on average than the others because the lineage is more rapidly lost by natural selection (or reverts to the other phenotype). We took all terminal branches where there is no change in the phenotype and analysed their lengths. This revealed significantly shorter branches in non-transformable strains (Ab: 90x shorter, p=1.14×10^-09^; Lp: p=0.03; Wilcoxon tests). Again, this suggests that non-transformable strains tend to be removed from the population by natural selection.

### Transformation impacts recombination rates and genetic linkage

Some of the proposed causes of selection for transformability, e.g., DNA as a source of nutrient or for repair, cannot be tested with our data. But the hypothesis that natural transformation facilitates adaptation by favouring allelic exchanges can be tested. We identified recombination tracts in the terminal branches of the species trees which covered 2628 persistent gene families in Ab and 2325 in Lp (see Methods). When considering the phylogeny, transformable strains exhibited a higher recombination rate than the non-transformable ones (phyloglm T17, Lp: p=1.29×10^-15^; Ab: p=1.36×10^-3^). Hence, we can identify the expected association between recombination rates and transformation, even if its effect is small and non-significant when phylogeny is not accounted for (Wilcoxon tests, all P>0.05). To better understand the relation between recombination and transformation, we analysed strains whose transformation phenotype changed recently. We also observed greater cumulated lengths of recombination tracts when the strain became transformable in the extant branch than when it became non-transformable, but the result was significant only in Ab (Figure 2A, Table S4). This suggests that the gain of transformation is responsible for an increase in recombination rates.

**Figure 2.**
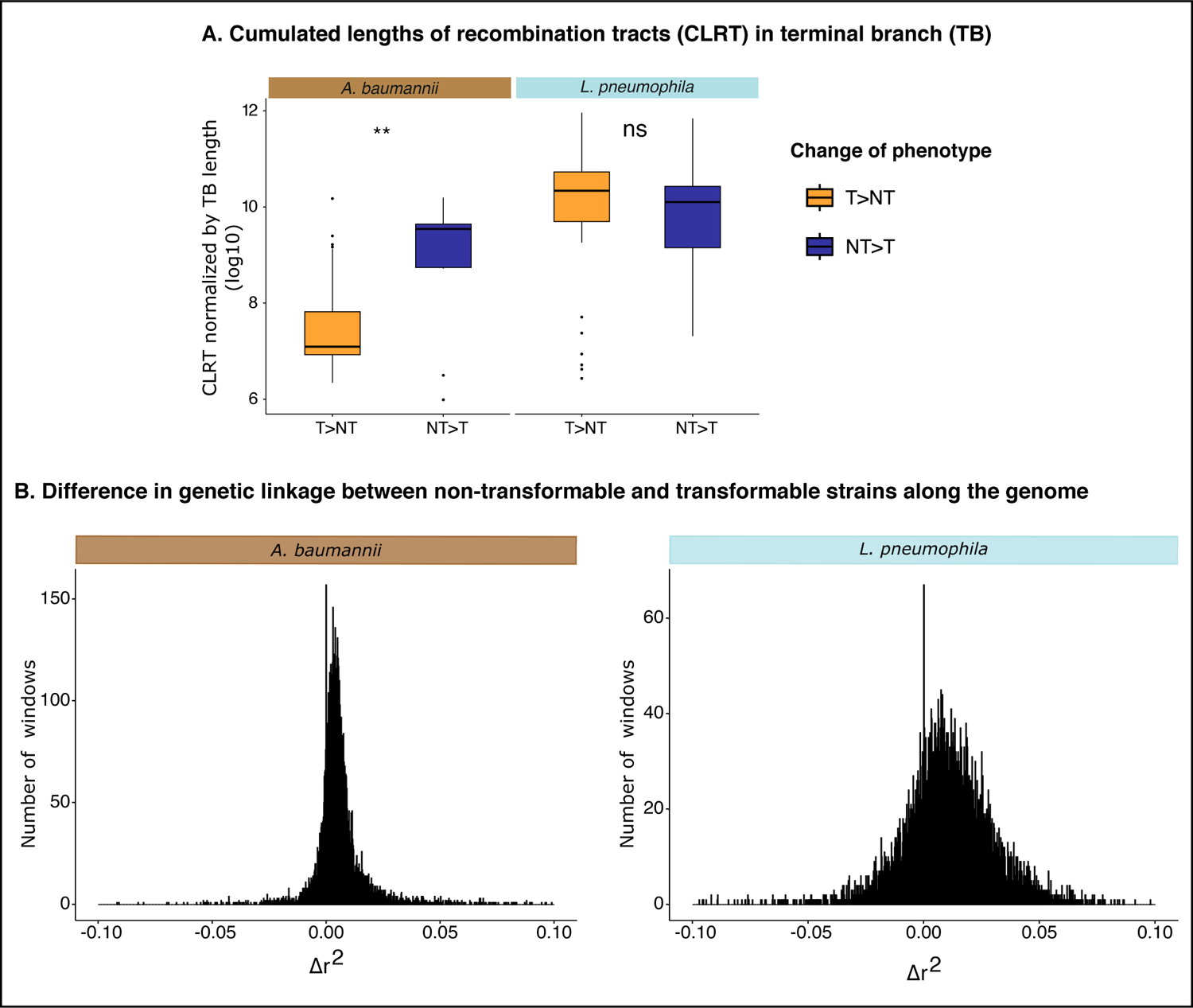
Association of the transformation phenotype on recombination. A. Distribution of cumulated length of recombination tracts (CLRT) (Log10-transformed) in terminal branches (TB) divided by the length of the latter in relation to the inference of changes in the phenotype. Wilcoxon test, *: p<0.01, **: p<0.01, ***: p<0.001, ****: p<0.0001 B. Distribution of Δr^2^ values in the [-0.1;0.1] interval computed in 500 nt screened windows where r^2^ was not null in both T and NT populations. The full span of the distributions is presented in Figure S4. Δr^2^ was calculated as r^2^mean(NT)-r^2^mean(T)

Since transformation favours allelic exchanges, it could have an important impact in breaking genetic linkage. This hypothesis can be tested by analysing the patterns of linkage disequilibrium. We calculated the squared correlation (*r*^2^) between bi-allelic values at two loci in windows of 500 nt along the genome of transformable and non-transformable strains. The difference between the two (Δr^2^) was significant (average 0.0092, Wilcoxon, p<2.2×10^-16^) (Figure 2B, S4), indicating higher correlation, hence higher linkage disequilibrium, in non-transformable strains. This agrees with the above hypothesis and suggests that transformation may facilitate breaking deleterious allele combinations or creating novel adaptive ones.

### Transformation rate variations rarely depend on its molecular pathway

To understand the causes of the loss of transformation we focused initially on the most obvious candidate genes: the ones directly involved in the molecular process (Figure 3). They are the most susceptible to influence the transformation phenotype. We made a survey of the literature to list genes associated with transformation and searched for their presence in every genome (Table S8). The gene *pilQ* was interrupted by a phage in one Ab strain and pseudogenized in three Lp (all four strains are non-transformable). In Ab some genes for minor pilins could not be identified in a few genomes of both transformable and non-transformable strains (PilEVX). Knock-out of minor pilins was enough to block transformation in Ab W068 [29]. Their absence in transformable strains may be explained by the minor pilins having a more diverging sequence than our search constraints would allow. Pilins evolve quickly by point mutation, HGT and duplication processes, complicating their detection in draft genomes. More importantly, the gene *comM,* encoding a helicase involved in the homologous recombination of transforming DNA into the chromosome, was often pseudogenized. This was significantly more frequent in non-transformable than in transformable strains (*X*^2^, p=1.46×10^-04^) (Figure 3) and in clinical strains relative to environmental ones (*X*^2^, p<2.2×10^-16^). The inactivation of this gene was most often caused by its interruption in the MgCh domain by the integration of MGEs. This has been described in clinical strains, where AbaR and AbGRI resistance islands integrate in this region. But contrary to previous analyses our dataset has only a small number of clinical strains. Our detailed analysis of this locus revealed AbaR islands in a fourth of *comM* inactivations. In the other cases, we found in these regions MGEs with a plethora of insertion sequences, integrons and defense systems. For example, the gene was interrupted by a locus encoding CBASS (39% of the times) and Zorya (10%). The latter interruption was only observed in environmental strains. This is consistent with the existence of genetic conflicts between MGEs and the host regarding natural transformation beyond the selection for the spread of antibiotic resistance genes in clinical strains.

**Figure 3.**
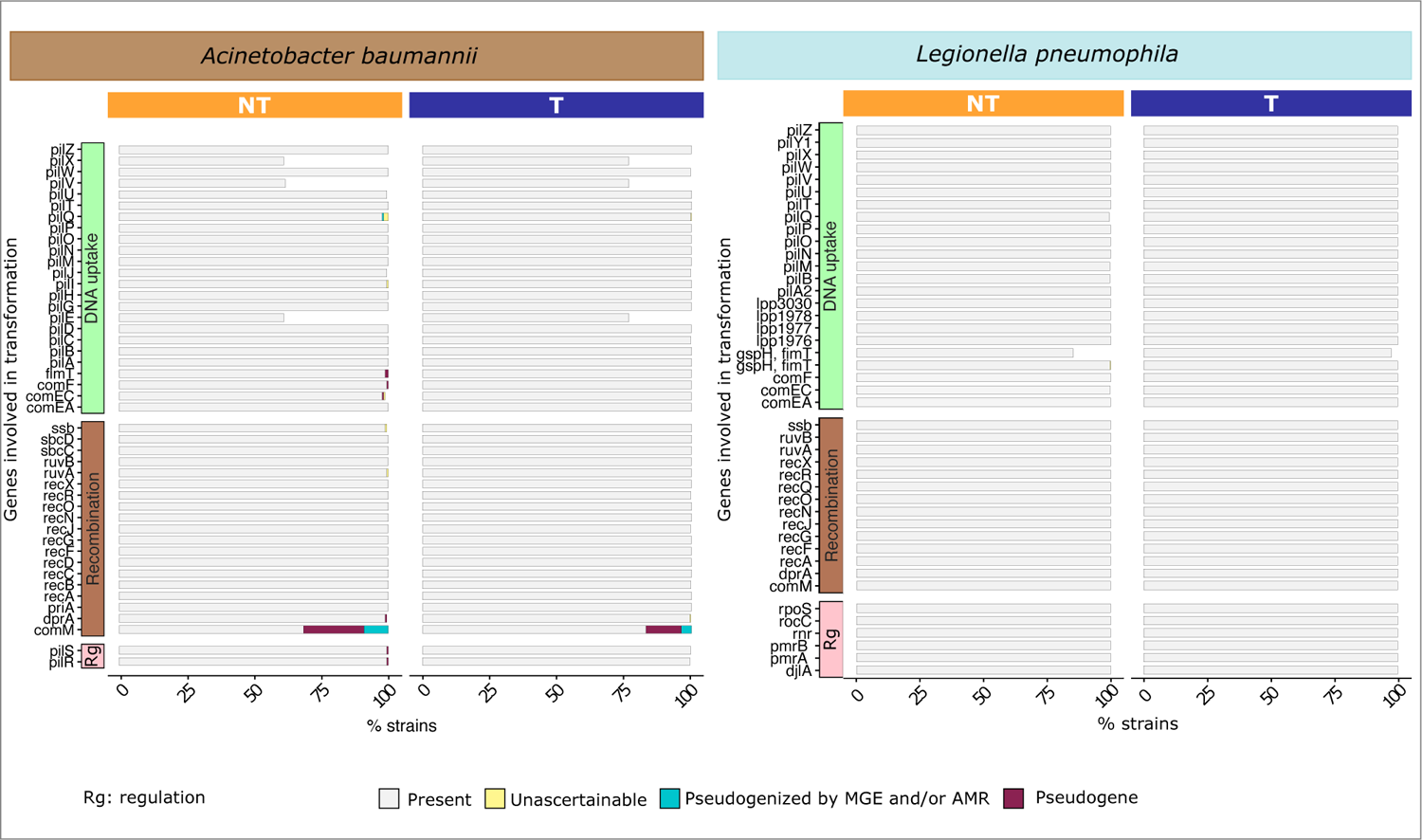
Presence of genes involved in natural transformation in *Acinetobacter baumannii* (left) and *Legionella pneumophila* (right). The genes were divided regarding their function: DNA uptake, recombination, and regulation (Rg) of transformation.

Interrupted transformation genes might be recovered by recombination and allow re-acquisition of the transformation phenotype. To test this possibility, we analysed the patterns of recombination on the locus in the 386 genomes of Ab with a complete *comM*. We found that *comM* was covered by a recent recombination tract in 35 samples. This raises the possibility that inactivated *comM* genes may become functional again by recombination with homologous DNA arriving from other strains. Yet, recombination salvaging comM does not happen more often than expected by chance (*X*^2^, p=0.72). This process is easier to achieve for genes like *comM* whose inactivation does not completely block recombination. With the exception of *comM* in Ab, variations in transformation rates were rarely explained by the inactivation of transformation-related genes.

### MGEs shape transformation rate variations

To identify the genetic determinants responsible for shifts between transformability and non-transformability, we performed a unitig-based Genome Wide Association Study (GWAS_U_) on the binary transformation phenotype (GWAS^bin^_U_). We used a Linear Mixed Model (LMM) to correct for population structure using the recombination-aware phylogenetic trees and applied a Benjamini-Hochberg adjustment for multiple tests at 0.05. The relevant genetic variants were mapped to the gene families. In Lp, 378 gene families including 1 sncRNA gene were associated with the inhibition of transformation and 271 were associated with increased transformability (Figure 4A, Table S5). In Ab, 836 gene families were associated with the inhibition of transformation and 426 with increased transformability (Figure 4A, Table S5). Among the genes directly involved in natural transformation only unitigs mapping *comEA*, *rnr*, *recQ* in Lp and *pilP*, *pilT* in Ab were associated with non-transformability. Since these genes are part of the persistent genome this suggests that some variants may lower transformation rates. Only the unitigs mapping *comM* were positively associated with transformability, a consequence of the abovementioned frequent inactivation of this gene. This confirms the impact of *comM* inactivations on transformation and suggests that natural sequence variants of the other genes may affect transformation rates. Overall, the genes directly implicated in the pathway of natural transformation are a very small fraction of all genes identified by the GWAS^bin^_U_.

**Figure 4.**
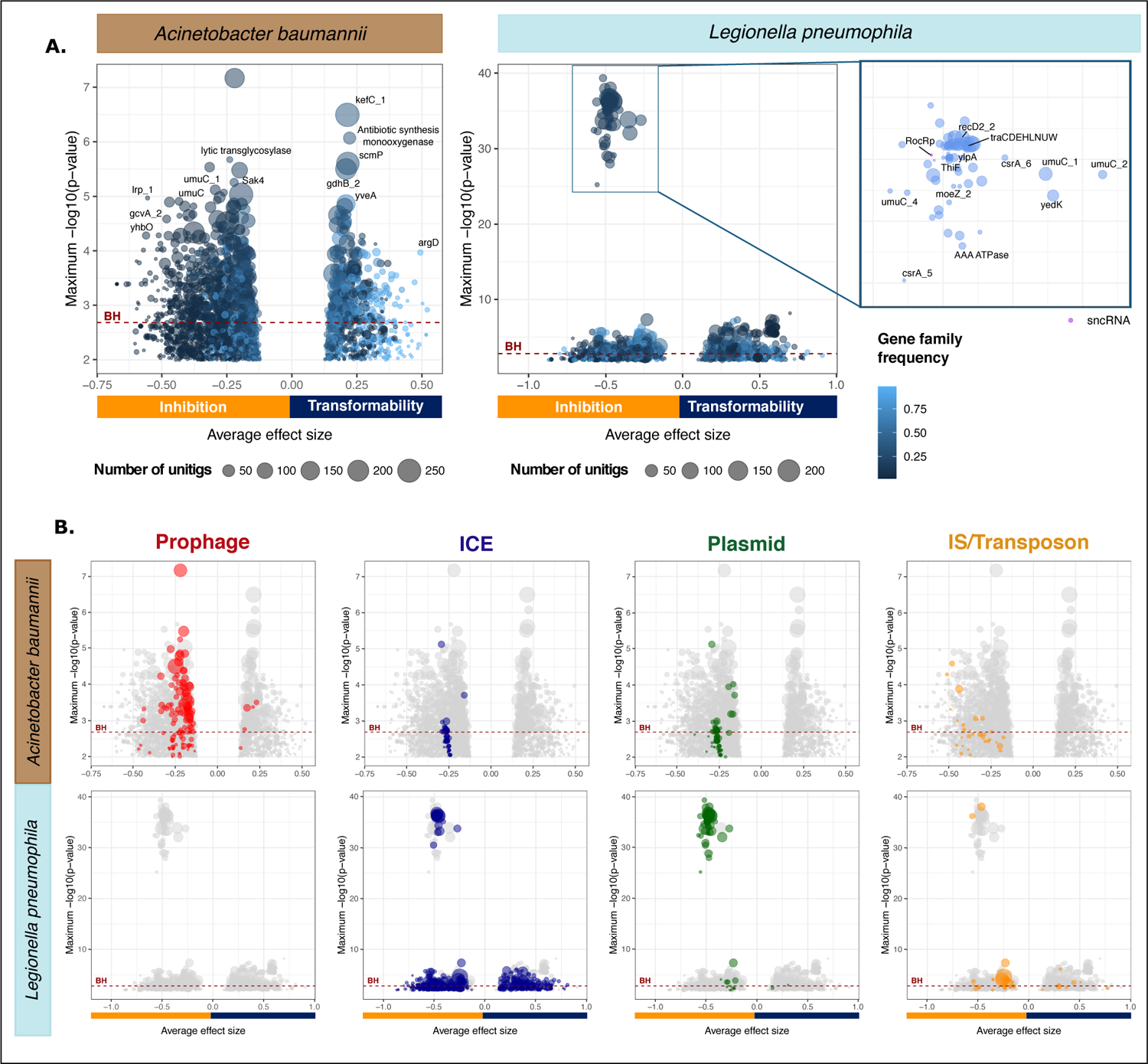
Volcano plots showing average effect sizes and significance of the association of the gene families with the transformation phenotype according to *GW AS^bin^_U_ Acinetobacter baumannii* and *Legionella pneumophila*. Each circle stands for a gene family. The size of the circle depends on the number of unitigs that mapped the gene in all the samples. The value on the x-axis corresponds to the average effect size of all the unitigs mapping the gene. The y-axis indicates how significant this effect can be by representing the maximal -log10-transformed p-value adjusted for population structure of all the unitigs of this gene. Significantly associated gene families are above the Benjamini-Hochberg (BH) threshold (red dashed line). The lower graphs (B) are similar to the ones on top (A), but gene families were colored in respect to their MGE.

Most of the strongest genetic determinants of transformation inhibition are in MGEs (in both species). We classed them in integrative conjugative elements (ICEs), plasmids, phages, and transposable elements (Figure 4B). Beyond the genes that were individually significantly associated with low transformation, many of the other MGE genes were collectively negatively associated with the loss of transformation, even when each individual effect was not significant (Figure 4B). For example, many Insertion Sequences, sometimes part of larger elements like plasmids and ICEs, are among transformation-inhibiting candidates in both Ab and Lp (Figure 4B).

Several prophage genes are negatively associated with transformation in Ab. Some of these functions can also be found in other elements, but others are very specific to phages as revealed by the analysis of their viral quotients (see Methods). Out of the 836 Ab transformation-inhibiting gene families from GWAS^bin^_U_, 24 matched proteins with very high viral quotient (higher than 0.9, see Methods). Among them, the most significantly transformation-inhibiting gene seems to be part of a prophage anti-defence system. We predicted its protein structure with AlphaFold and aligned it against a large dataset of protein structures using Foldseek. This analysis revealed structural homology to DarA, a protein that is involved in capsid morphogenesis and is a component of an anti-restriction system (found in *A. baumannii* ACICU, Figure S5) [30]. However, the homology is restricted to the N-terminal domain and this gene stands alone in the GWAS (the Dar system being composed of many proteins). Further work will be necessary to disentangle the function of this protein. In brief, the abundance of prophage genes in the GWAS suggests these elements have an important role in the inhibition of transformation in Ab.

We could not identify intact prophages in the Lp strains. Instead, the transformation-inhibiting candidates in this species were often associated with functions found on plasmids, including conjugation and the plasmid-encoded sncRNA RocRp (Figure 4). A little more than 15% of the Lp transformation-inhibiting genes identified in GWAS^bin^_U_(62/378) were carried by plasmids. It was previously shown that RocRp is plasmid-encoded and inhibits transformation [21]. None of the RocRp copies could be identified in the chromosomal contigs. To confirm that contig assignment was not affecting our results, we searched 113 complete Lp genomes from RefSeq where all 5 occurrences of RocRp were also in plasmids. Overall, 32% of the non-transformable strains encoded RocRp plasmids. This raised the following question: are the plasmid-associated genes in the GWAS^bin^_U_found because of their genetic linkage with rocRp or because they are independently associated with the inhibition of transformation? To answer this question, we built a linear mixed model where the presence of rocRp in the strain is a covariate of GWAS^bin^_U_ (GWAS^bin^_U_-cov). This model retained some significant associations between plasmid-associated genes and transformation in Lp (BH adjusted pvalue < 0.05, Figure S6), but the overall p-values of these genes were much smaller. In fact most of these genes were conjugation genes that also exist in ICEs suggesting their association might be retained because of some other transformation determinants that ICEs carry. Hence, the negative association between plasmids and transformation is largely driven by genetic linkage with the transformation-inhibiting sncRNA rocRp, albeit some other genes present in certain ICEs may be good candidates for secondary modulators of transformation rates.

Given the importance of MGEs in shaping transformation rates, anti-MGE defence systems could directly lower transformation rates by targeting incoming DNA [31] or indirectly increase transformation rates by preventing the acquisition of MGEs encoding transformation-inhibiting genes. We separated innate (e.g., restriction-modification systems) from adaptive (CRISPR-Cas) defense systems because the former may block MGEs and bacterial DNA arriving by transformation, whereas the latter are only expected to block MGEs (since they provide a specific defence). We detected a negative association between the number of putatively innate defence systems of a strain and its transformability in Lp and in Ab (phyloglm T12, Lp: p=1.03×10^-13^; Ab: p=3.61×10^-9^) and more specifically when the defense system was a restriction-modification system (phyloglm T13, Lp: p=4.34×10^-8^; Ab: p=1.30×10^-6^). In contrast, there was a positive association between the number of CRISPR-Cas systems carried by a strain and its transformability in both species (phyloglm T14, Lp: p=7.81×10^-9^; Ab: p=0.018). These results suggest that defense systems impact transformation rates in ways depending on their ability to specifically target MGEs.

### Transformation is associated with the loss of MGEs

The chromosome curing model suggests that genetic conflict between MGEs and the host arise because transformation cleans the bacterial chromosome from its MGEs. One would thus expect to find fewer MGEs in transformable strains than in the others. We analyzed the frequency of MGEs in relation to the transformation phenotype and the phylogenetic structure using a phylogenetic logistic regression [32]. Even though some prophages are negatively associated with transformation (see above), the number of prophages in Ab does not depend on the strain transformation phenotype (p>0.05; no prophages in Lp). When compared to the others, the chromosomes of transformable strains carry fewer Insertion Sequences (Figure 5; phyloglm T19, Lp: p=5.97×10^-5^; Ab: p=0.0016) and fewer conjugative systems (phyloglm T7, Lp: p<2.2×10^-16^; Ab: p=1.21×10^-7^). Of note, conjugative plasmids were significantly rarer in transformable strains in both species (phyloglm T9, Lp: p<2.2×10^-16^; Ab: p=1.38×10^-8^). Hence, many types of MGEs are less abundant in transformable strains.

**Figure 5.**
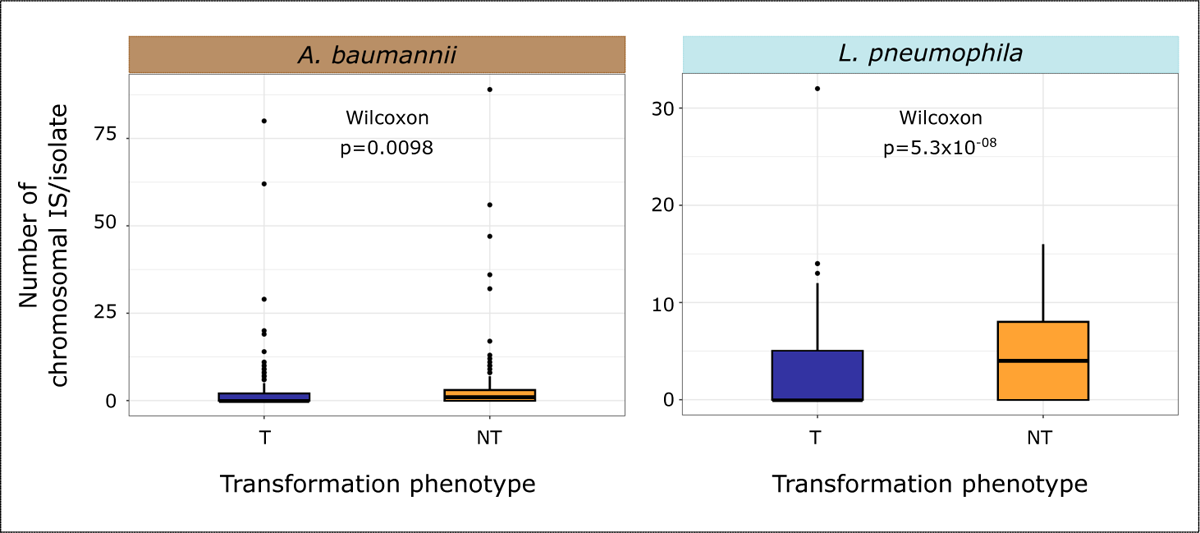
Distribution of the number of Insertion Sequences in the bacterial chromosome per isolate in transformable and non-transformable strains in *Acinetobacter baumannii* (left) and *Legionella pneumophila* (right).

If recombination cures the chromosome from MGEs, one expects to find an excess of homologous recombination targeting persistent genes that flank MGEs. For each persistent gene we used the number of times it was covered by a recombination tract in our collection as a proxy for its recombination rate. Based on the MGE flanking persistent genes previously identified in the whole collection, we were able to evaluate for each persistent gene if it had a MGE in its direct neighborhood in at least one genome. We observed that among persistent genes, those in the direct neighborhood of a MGE had higher recombination rates in Ab: conjugative systems (Ab:Wilcoxon, p<2.2×10^-16^), and insertion sequences (Ab: Wilcoxon, p<2.2×10^-16^), but not for phages (Ab:Wilcoxon, p=0.08). In Lp the association was significant when the MGE was an insertion sequence (Lp: Wilcoxon, p=1.7×10^-5^). Interestingly, recombination also targets at high frequency the persistent genes neighboring defense systems (Ab:Wilcoxon, p<2.2×10^-16^), which fits previous observations that these systems evolve rapidly and are often within MGEs [33]. In conclusion, core genes flanking MGEs show an excess of recombination tracts, as expected if transformation removes neighboring MGEs by recombination. These removals may become fixed in the population when they increase bacterial fitness.

## Discussion

We found wide within-species variations in transformation rates under similar growth conditions, with 64% of Ab and 52% of Lp being transformable beyond the detection limit of the method. Variations in transformation rates were previously shown in smaller samples of other species. In *A. actinomycetemcomitans* only 26% of strains are competent for transformation [34], and in *S. pneumoniae* and *H. influenzae* [17], [19] around two thirds of the strains are transformable. The ranges of variation in transformation observed in this study (4 orders of magnitude in Lp and 6 in Ab) are close to the ones of *H. influenzae* (6) and *S. pneumoniae* (4). They are probably underestimates because transformation rates may vary between environments (for which there is no data available). We found very little association between transformation rates and population structure, as previously observed in *S. pneumoniae* [17], but contrary to observations in *P. stutzeri* [34]. Instead of transformation rates following species phylogroups they are associated with the patterns of distribution of MGEs.

Allelic recombination decreases genetic linkage thereby rendering natural selection more efficient. Recombination tends to facilitate the fixation of adaptive mutations and alleviates the cost of deleterious ones [35], especially if it is fitness-associated [36]. It could thus be one of the key advantages of transformation. We found that transformable strains have slightly higher recombination rates than non-transformable ones. This effect is only identifiable when considering the effect of phylogeny, suggesting that recombination rates are poor proxies for transformation rates. This may be caused by the confounding effect of recombination mediated by MGEs. Strains with few MGEs could tend to recombine by natural transformation whereas those with many MGEs might tend to transform at low rates but still recombine as the result of conjugation or transduction. While natural transformation is expected to result in the recombination of persistent genes across the entire chromosome, MGEs will favour recombination of genes close to the regions where they integrate the genome [37], [38]. This could explain why there is a weak positive association between transformation and recombination rates, but a clear loss of genetic linkage associated with transformation: for similar rates of recombination, transformation provides DNA covering the core genome more uniformly thereby decreasing genetic linkage more efficiently than MGE-driven recombination that tends to favour the transfer of core genes close to the MGE integration site. To test this hypothesis, it will be necessary to disentangle in the future the contribution of different mechanisms of HGT to homologous recombination.

The simplest mechanistic explanation for the existence of many non-transformable strains is the inactivation of transformation-related genes by MGEs [13]. One key gene (*comM*) in one of the species (Ab) was often inactivated (22% of the strains) because of AbaR islands in clinical strains and many other diverse MGEs in the others. The extensive collection of Ab environmental strains we have used shows that *comM* is targeted by a much broader range of elements often encoding defense systems. ComM inactivation was also previously described in *H. influenzae* [39], *A. actinomycetemcomitans* [40] and several *Pasteurellaceae* [41], raising an interesting question: Why would ComM be specifically targeted for inactivation? The function of this protein was recently clarified. It is a helicase that facilitates homologous recombination between heterologous DNA [7]. Its loss reduces transformation by two orders of magnitude for the type of DNA tested in our assay (non-homologous segment flanked by homologous regions) [7]. In *V. cholerae* its absence has a very minor role in transformation with identical DNA [42]. This may explain why strains lacking ComM still have measurable rates of natural transformation. The lower recombination with heterologous DNA in *comM* mutants may have a small impact on the host fitness, since DNA uptake and recombination between homologous DNA may still take place. Yet, it may diminish the ability of the bacterium to remove MGEs from the genome (because distant strains with heterologous DNA are more likely to lack the MGE and lead to its deletion from the genome by transformation). A key question that has not been addressed before concerns the fate of the lineages lacking a functional *comM*. Beyond *comM* (and only in Ab), inactivation of competence genes rarely explains the observed variations in transformation rates, possibly because most of these genes also contribute to other important processes, such as DNA repair (recombination), adhesion, and virulence (type IV pilus) [43]. Hence, their inactivation would decrease the host and the MGE fitness.

What justifies the high variation in transformation rates across the strains and especially the observed frequent non-transformability? Many of our results are compatible with the idea that intragenomic conflicts between MGEs and the bacterial host govern the variation in transformation rates. We confirmed the negative association between transformation rates and the presence of RocRp-encoding plasmids in Lp [21] and *comM*-inactivating MGEs islands in Ab (known for AbaR) [44]. We showed that transformation rates vary following a jump process. Both observations are consistent with a negative impact of MGEs on transformation rates since these elements are frequently gained and lost [45]. Accordingly, we found that MGEs such as ISs, ICEs, plasmids, and phage (depending on the species) are systematically associated with low rates of transformation. For some types of elements, the significance of each individual gene family was low, e.g., for transposable elements, but grouping these elements per type confirmed they are associated with low transformation rates. Finally, intragenomic conflicts explain our seemingly contradictory observations that transformation rates are often below the detection limit even though these strains seem to be counter-selected.

The intragenomic conflict between MGEs and bacteria has been explained by the chromosome curing hypothesis [13]. In this model, bacteria use transformation to delete MGEs from chromosomes by recombination at flanking persistent genes and MGEs strive to counteract this mechanism. This is consistent with most of our observations described above. It is also consistent with our observation of higher rates of recombination on core genes flanking MGEs and lower frequency of MGEs in transformable bacteria. However, this does not explain why conjugative plasmids in Lp are repressing transformation via RocRp. Plasmids cannot be deleted from the chromosome because they are extra-chromosomal. They also usually lack persistent genes and transformation is not expected to delete them. In other bacterial species RocRp can be found in chromosomal islands and in ICEs [46]. Yet, in Lp this gene is exclusively found in plasmids where it has a very strong negative effect on natural transformation. This suggests that intragenomic conflicts between MGEs and bacteria regarding transformation extend to extra-chromosomal elements. We speculate that plasmids might block transformation to prevent the acquisition of incompatible plasmids, exploitative MGEs, or anti-plasmid defense systems. In any case, these results strongly suggest that intragenomic conflicts are not restricted to the effect of chromosome curing.

The interplay between anti-MGE systems and transformation is complex in the presence of MGE-driven intragenomic conflicts. These defense systems may decrease transformation rates by blocking the entry of exogenous DNA or they may increase them by blocking the acquisition of transformation-inhibiting MGEs. When we excluded the known adaptive defense systems (CRISPR-Cas) we found a negative association between defense systems and transformation rates. Many of these are restriction modification systems, by far the most abundant defense systems of Bacteria [47]. Restriction systems were shown to reduce transformation efficiency in specific strains of *P. stutzeri* [48], *H. pylori* [49], *Neisseria meningitidis* [8], *C. jejuni* [50], [51], *S. pyogenes* [52], and Ab [31]. CRISPR-Cas are adaptive immune systems and their presence is positively correlated with transformation in both species. The opposed associations of CRISPR-Cas and restriction-modification systems can be explained by the way they work. CRISPR-Cas systems cannot efficiently protect from transformation because, contrary to R-M, they cannot block generic heterologous DNA but only elements carrying the sequences matching their spacers. Hence, they have probably very little if any role in preventing the acquisition of homologous DNA by transformation. Since they protect the bacterium from the acquisition of MGEs that might block transformation, they have a net positive effect on transformation rates. This means that the impact of defense systems on the variation of transformation rates depends crucially on the specificity and adaptability of their mechanisms of action.

We found striking parallels between Ab and Lp, despite their very different lifestyles and gene repertoires. Both have widely variable transformation rates evolving according to a jump process and decreasing genetic linkage. Non-transformability is scattered across the species trees, seems to be counter-selected, and is associated with the presence of MGEs. This suggests common reasons for the variability of transformation rates. Intragenomic conflicts driven by MGE are compatible with most of the data, suggesting that bacteria-MGE interactions are key drivers of the evolution of natural transformation rates. If intragenomic conflicts seem to have a key role in the evolution of transformation rates, their underlying causes, e.g., chromosome curing of MGEs, are not necessarily the only reasons for the selection for natural transformation. Other putative advantages of transformation, gene transfer, nutrient acquisition, or DNA repair, are lost if the mechanism is inhibited by MGEs. These advantages have the potential to further increase the intragenomic conflict between the bacterium, which benefits from natural transformation, and the MGEs that benefit from blocking it.

## Materials and Methods

### Bacterial strains, origin and typing

We analyzed draft assemblies of 830 *Lp* clinical strains from the Centre National de Référence pour *Legionella* (CNRL) collection and of 510 *Ab* environmental and clinical strains, all listed in Table S6. We sequence typed Ab collection with *mlst v.2.19.0 (*https://github.com/tseemann/mlst). The Lp collection was assembled from clinical isolates collected in France from 2018 to 2020 and was sequence typed by the CNRL. This publication made use of the PubMLST website (https://pubmlst.org/) developed by Keith Jolley [53] at the University of Oxford. We also typed *Ab* capsules with *Kaptive v.2.0.3* [54], [55]. The search for antimicrobial resistance genes was done with ABRicate (https://github.com/tseemann/abricate) which used the following databases ARG-ANNOT [56], CARD [57], MEGARes [58] and VFDB [59].

### Plasmid constructions

Plasmid pJET.Lp-*pilMNOPQ::nLuc* was constructed by cloning the nanoluc gene along with flanking arms of 2000 bp from the *pilMNOPQ* locus obtained from the genomic DNA of *Legionella pneumophila* strain Paris. Plasmid pJET.Ab-*pilMNOPQ::nLuc* was constructed by cloning the nanoluc gene along with flanking arms of 2000 bp from the *pilMNOPQ* locus obtained from the genomic DNA of *Acinetobacter baumannii* strain A118.

### Construction of pangenomes

We removed poor quality draft assemblies with *PanACoTA v.1.2.0* [60] keeping drafts with less than 100 contigs when the sum of the 100 largest contigs was at least 90% of the genome (L90≤100). We excluded strains that were too similar to already included strains or too distant from the other strains in terms of mash distance to be part of the species (Lp= [10^-6^;0.1], Ab= [10^-6^;0.06]). After filtering, we were left with 786 Lp isolates and 496 Ab isolates.

We annotated the draft assemblies (*prokka* from *PanACoTA v.1.2.0 completed with eggnog-mapper 2.1.9*) and built the pangenome of each collection using single-linkage clustering to form families of proteins with at least 80% identity (using *mmseqs2 v.12-113e3* within *PanACoTA v.1.2.0*). We defined the persistent genomes of each species. A pangenome family was considered persistent if at least 95% of the genomes had a unique member of this family (Lp: 11932 pangenome families and 2326 strict-persistent pangenome families, Ab: 31103 pangenome families and 2629 strict-persistent pangenome families).

### Natural transformability assay for L. pneumophila

*L. pneumophila* strains from frozen stock cultures were thawed, 5 µl of the cells were spotted onto CYE solid medium plates and incubated for 3 days at 37°C. Cells from the plates were subsequently resuspended into 100 µl liquid AYE medium in 96-well plates using a 96-well Scienceware® replicator and allowed to grow for 3 days at 37°C in a shaking incubator. Next, 2 µl of this culture was transferred to 100 µl of fresh AYE medium in 96-well plates containing 20 ng/µl of pJET.Lp-*pilMNOPQ::nLuc* plasmid DNA and allowed to grow for 3 days at 30°C in a shaking incubator. As this plasmid is non-replicative in *L. pneumophila,* DNA molecules which are internalized undergo a double recombination event allowing the insertion of *nanoluc* gene in the *pilMNOPQ* locus and subsequent expression of the NanoLuc® Luciferase enzyme. Subsequently, 80 µl of cells were mixed with 20 µl of the Nano-Glo® Luciferase Assay Substrate and Nano-Glo® Luciferase Assay Buffer followed by incubation at room temperature for 10 minutes. The expressed NanoLuc® Luciferase enzyme was reported as luminescence units (LU) on a Promega GloMax® Navigator plate reader. The optical density of the cell suspension at 600 nm was detected on a Tecan plate reader. Relative luminescence units (RLU) were calculated by dividing the luminescence values by the optical density values. RLU values were used as a proxy for transformability of the strains. The assay was repeated independently for each strain in the collection between 3 to 6 times.

### Natural transformability assay for A. baumannii

Natural transformation in *A. baumannii* requires induction by agarose. Agarose soluble extract media (ASEM) was prepared by adding 2 g of Agarose D3 (Euromedex) to a 5 g/L solution of Tryptone media (Bacto™). This suspension was vortexed for 5 minutes and subsequently centrifuged to sediment the insoluble agarose particles. The supernatant was collected and filtered using a 0.22 µm filter. *A. baumannii* strains from frozen stock cultures were thawed, subsequently 2 µl of cells were transferred to LB medium (Lennox formulation) and incubated overnight at 37°C. The following day, 2 µl of cells were transferred to ASEM containing 2 ng/µl of pJET.Ab-*pilMNOPQ::nLuc* plasmid DNA using a 96-well Scienceware® replicator and allowed to grow overnight at 37°C. As this plasmid is non-replicative in *A. baumannii,* DNA molecules which are internalized undergo a double recombination event allowing the insertion of *nanoluc* gene in the *pilMNOPQ* locus and subsequent expression of the NanoLuc® Luciferase enzyme. Subsequently, RLU values were calculated similarly as described above for *L. pneumophila*. The assay was repeated independently for each strain in the collection between 4 to 6 times.

### Assessment of the transformability phenotype

The number of Ab or Lp transformants is linearly correlated with the luminescence signal. Each strain transformation rate was then calculated as the average of the log10-transformed OD-corrected luminescence value on all the replicates since the assay gave reproducible results between replicates: Spearman correlation coefficients were on average 0.72 in Lp and 0.90 in Ab (Table S7). The log10-transformation of the OD-corrected raw luminescence values reduced the skewness of the different replicates and allowed us to compare them. The threshold between transformable and non-transformable strains (Lp: LumR/OD=300, Ab: LumR/OD=400) was set based on the maximum of the transformation rates of a non-transformable strain: the Lens strain for Lp (120 replicates) and an engineered Δ*comEC* strain for Ab (28 replicates).

### Competence genes presence/pseudogenization

We gathered from the literature a list of competence genes and genetic elements (protein coding genes, sncRNA) involved in the transformation process (Table S8). We checked in all our strains for the presence of these elements. We retrieved their protein sequences from Paris and Philadelphia strains for *Lp* and from *A. pittii* PHEA-2 and *A. baumannii* D1279779 for *Ab* respectively. We searched for homologous regions to proteins involved in transformation using *tblastn v.2.12.0* (-evalue 0.001, -seg no) in the genomes in our collection [61]. We deemed a gene *present* if the alignment given by tblastn had more than 80% identity with the query protein and covered more than 80% of it. sncRNA genes were searched with *blastn v.2.12.0* (-evalue 1e-10). A sncRNA gene was present if the alignment with the query gene had 100% identity and covered more than 90% of it.

When the protein coding gene was not present, we searched for pseudogenes. We considered that we identified a *pseudogene* of a competence gene when we identified a gene/pseudogene more than 80% identical to the query protein and with an alignment covering between 20 and 80% of the protein (using the output of tblastn as described above). Some genes may look like disrupted simply because they are at the border of a contig. To account for contig borders, we considered that if in addition the alignment was located at less than 50 nucleotides from the end of the contig, we could not classify it as present, missing or pseudogene and called them *unascertainable*. Some genes marked as pseudogenes are split on two different contigs. They may be interrupted or not in the actual genome. If both alignments are at the border of the contigs (less than 50 nucleotides from the end of the contig) this is consistent with the two hypotheses, and we marked them as *unascertainable*. If they are not both at the border, we marked them as *pseudogenes*.

Pseudogenes with large nucleotide insertions were further characterized to assess if the disruption was due to MGE and/or AMR genes. A supplementary step to assess the nature of the interruption was necessary when these pseudogenes were split on two contigs. We took the two alignments of the gene (one in each contig) and ordered them in relation to the known full gene sequence. This allowed to identify the sequence interrupting the gene. This was only the case for ComM protein, which has 3 domains organized in the following order: chlI (PF13541.9), Mgch (PF01078.24) and MgchC (PF13335.9). We relied on the coordinates and order of the domains in our alignments to identify the correct region of interruption in which we would search for MGEs and AMR genes.

In *Acinetobacter baumannii*, we had to devise a specific method to search for pilA because of the high variability of its sequence across the species [62]. We searched the most conserved region of pilA, the pilin domain (PF00114) in the representative sequence of each gene family with *hmmsearch (hmmer/3.3.2).* We considered that the pilin was present when the sequence score of the alignment was above the gathering threshold score cutoff (--cut_ga). Every gene family (34) presenting this domain was considered encoding PilA and every strain having this gene family was deemed having *pilA*. We checked if the strains lacking these gene families could actually have an even more diverging sequence for *pilA.* We searched for homologous regions of four PilA proteins (QNT88830.1, WP_000993715.1, WP_031953428.1, WP_000993729.1) previously studied in *A. baumannii* [62] using *tblastn v.2.12.0* (-evalue 0.001, -seg no) in the genomes of those strains. We relaxed the constraints compared to other genes and considered the gene *present* if the alignment with the query had more than 40% identity, 20% coverage and a total length between 100 and 200 amino-acids.

We also searched for the presence of the rocRp gene in the complete genomes of Lp from RefSeq downloaded on May 2023 using *blastn v.2.12.0*. We deemed it present when the same criteria as above were met (100% identity, 90% coverage). To check that chromosomes do lack rocRp, we made a complementary analysis using a lower threshold of identity (90%). This analysis also failed to reveal chromosomal versions of the gene.

### Variant calling and estimation of genetic linkage

We identified single nucleotide polymorphisms (SNPs) and small insertions and deletion (indels) in our strains using as a reference genome the Paris strain for *Lp* and the AB5075 strain for *Ab*. The identification was done using *snippy v.4.6.0 (*https://github.com/tseemann/snippy). We annotated them and predicted their functional effects with *snpEff v.4.3* [63].

From the previous variant calling we only kept biallelic SNPs that were polymorphic in transformable strains and in non-transformable ones. We defined non-overlapping windows of 500 bp that scanned the whole genome. We considered pairs of SNPs (A,B). In each pair, we calculated p_r_ the frequency of the reference allele of SNP A (A_ref_) and q_r_ the frequency of the reference allele of SNP B (B_ref_). We also calculated x_rr_ the allelic frequencies of each pair of SNPs (A_ref_, B_ref_). We were then able to compute the D measure of linkage disequilibrium, its normalized value D’ and the r^2^, the square of the correlation coefficient of each pair of SNPs as follows [64].

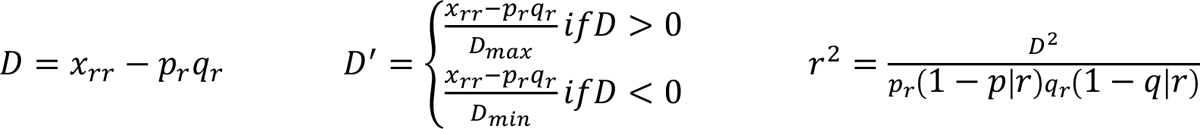

with

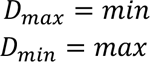

We compared the distribution of r^2^ in each genomic window between transformable and non-transformable populations. To assess if there was any difference in genetic linkage along the genome between transformable and non-transformable strains, we computed an average r^2^ for each window in both populations and tested with a Wilcoxon rank-sum test if the difference between the two, Δr^2^, was positive, i.e., if we rejected the null hypothesis H_0_, Δr^2^≤0.

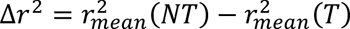

### Phylogenetic inference and analysis of recombination

We took the gene families that were regarded as persistent in Ab and Lp. In each species, we aligned each gene family with *mafft v.7.467* [65] within *PanACoTA v.1.2.0* (default parameters) [60] at the protein level. We back translated protein alignments to nucleotide ones (i.e., we replaced each amino acid by its original codon) because the latter provide more signal when one studies polymorphisms at the species level. Alignments at nucleotide level were concatenated to make two matrices of alignments of 2629 strict-persistent genes in Ab and 2325 strict-persistent genes in Lp ordered according to the persistent genes order and orientation of a complete genome of the collection (Lp: Paris strain; Ab: AB5075 strain). If a persistent gene was missing from the complete genome, its position and orientation was inferred from the most often frequent position and orientation it had in the collection. These matrices were then used as an input to *IQTree v.1.6.12 modelfinder* [66]–[68] to build the phylogenetic tree of each species.

We made a subsequent step of phylogenetic inference to account for the presence of recombination. In Ab in addition to the alignment described before, we also generated an alignment in which the alignments at the nucleotide level of the persistent genes were concatenated in a random order so as to see to what extent the correctness of the phylogeny and the recombination inference could impact the results (Figure S7). To account for the presence of recombination, we took the previous tree and used it as a starting point for *Gubbins v.2.4.1* [69]. *Gubbins v.2.4.1* allowed us to mask the regions in the alignment whose polymorphism was due to recombination. We finally built the phylogenetic trees based on the recombination region-free alignment with *IQTree v.1.6.12 modelfinder* according to which the best-fit model based on BIC was TVM+F+I+G4. To ensure of the branch robustness, we performed a 1000 ultrafast bootstrap.

Trees were rooted based on outgroups: *L. longbeachae* for *L. pneumophila* and *A. baylyi ADP1* for *A. baumannii*. We added each outgroup to their respective collection. We built their pangenomes, defined their strict-persistent genomes and aligned them in the same manner we did for the initial collections and with the same parameters. We finally built the phylogenetic tree based on this alignment with *IQTree v.1.6.12 modelfinder* (best-fit model: TVM+F+I+G4; 1000 ultrafast bootstraps) so as to determine the position of the root in the recombination-free phylogenetic tree. All trees are listed in Table S1.

We compared the changes in topology and branch lengths between the recombination-unaware and the recombination-free phylogenetic tree by measuring their weighted Robinson-Fould (wRF) metric (*phangorn* R package: *wRF.dist* function). This metric counts the minimum number of branch rearrangements needed to transform one tree into another and weights each branch by its length. The average root-to-tip distance of the phylogenetic tree was calculated by averaging the distance to the root of each tip (*adephylo* R package: *distRoot* function).

We also used the information on recombination tracts covering the persistent genes to assess the likelihood that neighboring non-persistent genes arose or were affected by recombination.

### Phylogenetic signal

We looked for phylogenetic inertia of the transformation trait in the rooted recombination-free phylogenetic trees. We calculated the phylogenetic signal on the log10-transformed transformation rates with Pagel’s λ and Blomberg’s K (*phytools* R package: *phylosig* function with test=TRUE to conduct a hypothesis test of K or λ) [70] and on the binary transformation phenotype with Fritz and Purvis’ D statistic (*caper* R package: *phylo.d* function and its default parameters) [71].

### Model of evolution for the trait

We assessed the models of the quantitative evolution of the trait. For this we used the log10-transformed transformation rates across the rooted recombination-free phylogenetic trees. We tested all the models mentioned in Table S3 for the trait evolution with *fitContinuous* function from *geiger* R package [72] and with *fit_reml_levy* function from *pulsR* R package [27]. We used the AIC as the selective criterion for the quality of model fit.

### Ancestral reconstruction of the transformation trait

We reconstructed the ancestral states of the binary transformation trait along the recombination-free rooted phylogenetic tree using the MAP prediction method and the F81 evolutionary model of *PastML v.1.9.34* [73]. We only kept nodes for which one of the possible reconstructed states had a marginal posterior probability superior to 0.6. We then assigned to the node the transformation state with the highest marginal posterior probability.

### Plasmid identification in draft assemblies

We classified contigs as plasmid by calculating the weighted gene repertoire relatedness (wGRR) of each contig against each RefSeq plasmid genome. We first searched for sequence similarity between all of their proteins using blastp (*blast+ v.2.12.0*). For each pair of contig/plasmid, the wGRR takes into account their number of bidirectional best hits and their sequence identities and gives an assessment of their gene repertoires similarity with the following formula.

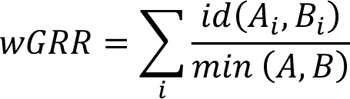

For a pair of elements A and B, id(A_i_, B_i_) is the sequence identity of the i-th pair of homologous proteins. Min(A,B) is the number of proteins in the smaller element. The wGRR of two elements is thus defined as the sum of the identity for all pairs of homologous proteins normalized to the number of proteins found in the smallest element.

Contigs with a wGRR superior to 0.9 were kept and preliminarily assigned to the corresponding plasmid. These contigs could either be from a plasmid or be a part of the chromosome carrying a MGE similar to the plasmid. Contigs respecting the wGRR criterion, shorter than 500 kb (the maximum size of Ab and Lp RefSeq plasmids being around 300kb) and carrying less than 50 % of persistent genes were deemed part of a plasmid (Figure S8). In Lp (Ab), 97.4% (43.3%) of the plasmids had no persistent gene at all. If the total length of the contigs assigned to the plasmid in a sample was longer than 40% of the actual plasmid length, we took this as indication of the presence of the whole plasmid in the sample. Contigs longer than 500 kb and with at least twice the number of proteins than the plasmid it was assigned to were deemed to be part of the bacterial chromosome.

We encountered in Ab two plasmids, Ab TG29392 plasmid pTG29392_1 from Ab TG29392 strain and plasmid pVB2107_2 from Ab VB2107 strain, which were completely integrated into chromosomal contigs (hits covering more than 50% of the plasmid). pTG29392_1 was integrated in one contig in 14% of the samples and pVB2107_2 in one contig in 3% of the samples. We classed these contigs as chromosomal contigs.

### Identification of MGEs and defense systems

We characterized the different types of mobile genetic elements. Conjugative elements were searched using *Macsyfinder 20221213.dev* [74], [75] and the *CONJScan/Chromosome* model (https://github.com/macsy-models). We detected conjugative systems (CONJ) which are complete conjugative systems formed by a T4SS, a type IV coupling protein (t4cp) and a relaxase, mobilizable systems (MOB) which are systems that have a relaxase that is either alone or co-localizing with a CONJ component but not enough of the latter to make a functional conjugative system. We checked if two contigs of a sample presented complementary incomplete CONJ on their extremities. If so, we included them with the other conjugative systems.

Available methods cannot precisely delimit the Integrated Conjugative Elements (ICEs). Hence, we defined them as regions at a distance of less than 10 kb from a conjugative system on the contigs that we did not identify as plasmid-like (cf. Plasmid identification in draft assemblies). This is a conservative estimate, given the average size of ICEs (typically more than 40kb for those of Proteobacteria, [76]). Insertion sequences were searched using *ISEScan v.1.7.2.3* [77]. We used the option --remove-ShortIS to remove incomplete IS elements that is to say IS elements of length inferior to 400 bp or single copy IS element without perfect terminal inverted repeat.

Putative prophages were initially searched using *VirSorter v.2.2.3* (docker://jiarong/virsorter:latest) [78]. We further analyzed the resulting elements with *CheckV v.0.7.0* [79] and kept the ones deemed of high and medium-quality. These putative viral regions were then annotated using *PHROG v.4* [80]. We built the hmm profiles of each PHROG based on their multiple alignments with *hmmbuild* (*hmmer/3.3.2* default parameters). We searched for those profiles in the previous viral regions with *hmmsearch* (*hmmer/3.3.2* default parameters). We also calculated a viral quotient for each PHROG annotation with the following formula:

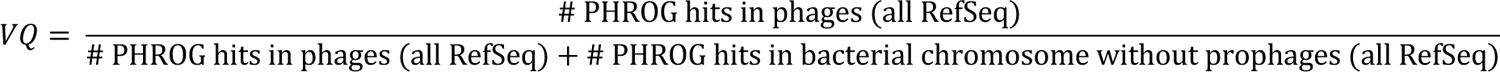

Gene families with a viral quotient superior to 0.9 were specific to phages and were as such labelled as phage-specific genes. The final list of prophages was obtained from the preliminary list by including information on the core genes (that should not be within prophages) and on the PHROG hits (which should be in prophages). Each putative prophage comprised between two persistent genes in a putative prophage containing at least 5 genes among which one had a PHROG annotation and whose average viral quotient is more than 0.5 was deemed to be a prophage. Integrons were searched for using *IntegronFinder v.2.0.2* [81]–[84].

We also characterized defense systems. CRISPR-Cas systems were searched using *CRISPRCasFinder v. 4.2.20* [74], [75], [85], [86]. We retained the ones verifying the following criteria: the presence of a Cas gene in the corresponding genome, the presence of more than 2 spacers in the array, and imposing a mean size for spacers of at least 15 bp. We identified the targets of these spacers by running *blastn* and its task option of *blastn-short* (*blast+ v.2.12.0*) on our collection of genomes, available RefSeq plasmids and GenBank phages setting the same parameters as *CRISPRTarget* (*CRISPRSuite*) [87]. We searched for other potential defense systems with the *DefenseFinder* models v.0.0.3 [47] using *Macsyfinder 20221213.dev* [75]. This version searched for 109 defense systems.

### Presence of MGEs in recombination regions

The analysis of *Gubbins* to detect recombination was made on the persistent genes (present in most genomes, see above). To know if recombined regions potentially encompass MGEs (which are never persistent genes), we had to devise a different method. First, we counted for each persistent gene how many times it was part of a recombination tract, thus obtaining a measure of their recombination rate. In parallel, we identified the closest upstream and downstream persistent genes for every characterized MGE. Of note, we could not determine the flanking persistent genes of 39.1% (30.0%) of the MGEs because none was present in one of the directions in the contig. Among these, 9.5% (24.2%) were on contigs classified as plasmids in Ab (Lp). These elements were excluded from the analyses of genomic neighborhood. We then separated the persistent genes flanking MGEs from the others. Finally, we compared the frequency of recombination of persistent genes of these two groups using a Wilcoxon test. We classed the loci of MGE integration into several categories: conjugative systems, phages, ISs and defense systems hotspots.

### Genome Wide Association Study

We performed a genome-wide association study (GWAS) to identify genetic determinants of transformation. We used a unitig-based approach (GWAS_U_). A unitig is an unambiguous combination of k-mers that represents the sequences without branch in the assembly graph. This approach allows to encompass information: on SNPs and large and small indels. We can thus test all these different levels of genetic variants all at once in the GWAS_U_. This was done on the binary transformation phenotype (TF^bin^) using respectively 786 *L. pneumophila* genomes and 496 *A. baumannii* genomes with *pyseer v.1.3.9* [88]. The unitigs were called on our collection of draft assemblies with *unitig-caller* (from conda installation of *pyseer*). The association between the unitig or the gene presence/absence and the transformation phenotype was assessed with a linear mixed model (LMM). The recombination-free phylogenetic tree allowed us to generate a distance and a kinship matrix using the scripts coming with *pyseer*, which were used in the linear-mixed model to control for population structure. To address the problem of multiple comparisons in our analysis, we applied a Benjamini-Hochberg adjustment on the p-value of association already adjusted for population structure [89]. We deemed the association between the presence of a unitig (or a gene) and the transformation phenotype significant when the adjusted p-value was inferior to 0.05.

To study the functions of genes with unitigs that were deemed significantly associated with changes in transformation rates, we first identified the genes having the unitigs with the *annotate_hits_pyseer.py* script provided by *pyseer*. To get a clearer picture of the associations, we summarized the characteristics of each gene association over all the unitigs mapping the gene; notably its statistical significance and its positive or negative effect on the phenotype. We only consider genes that are mapped by at least one significant unitig. These genes can be mapped by unitigs with significant effect of different signs (positively and negatively associated with transformation), in which case we exclude them from the main analysis and study them separately (see below). We are then left with genes mapped by unitigs (often many) for which those significantly associated with transformation are all of the same sign.

Following *pyseer summarize_annotations.py*, we summarized the positive or negative association between these gene families containing the unitigs and the phenotype, its effect size (by averaging it over all the unitigs of the significant effect sign that mapped the gene), its significance (by assigning the p-value of the most-significantly associated unitig with the gene), its minimum allele frequency and the number of unitigs mapping them. We added on top of the functional annotation the information on the presence of the gene family in a MGE or a secretion system (as defined above). In the case of *Lp*, we also studied the unitigs in the light of the presence of sncRNA.

### Detection of genes with variants of opposite effect on transformation

Some gene families were associated in an ambiguous way with the transformation phenotype. In Ab (Lp) 12% (43%) of the genes with one unitig significantly associated with variation in transformation rates also had unitigs whose effect was significant and of opposite sign. We treated those genes separately. Indeed, such a situation could be explained by unitigs carrying variants of the same gene (or same type of MGE) differently associated with the phenotype. So, averaging the effect size as we did before over all the unitigs of the gene would give misleading information on the association (e.g. if two unitigs have effects of symmetrical amplitude the average effect is zero, like for genes lacking unitigs altogether). We mapped the unitigs on the genes and called the variants with *snippy v.4.6.0 (*https://github.com/tseemann/snippy). We reduced the regions of interest to the regions that overlapped between unitigs of opposite association when this was possible. We summarized the positive or negative association to the phenotype, the effect size, the significance, the minimum allele frequency and the number of unitigs mapping to each variant. This procedure identified 52% (out of the 12%) of the genes in Ab and 66% (out of the 43%) in Lp. These genes with overlapping unitigs of opposite sense will be candidates for future analyses. Given their abundance, we further enquired on the regions with such unitigs in Lp. In Lp, half of them were in gene families of ICEs.

### Strains specificities affecting/affected by transformation rates

To characterize the association between traits across the tree, we need to take into account the phylogenetic structure of the bacterial population. We used phylogenetic logistic regression (*phyloglm* function from *phylolm* R package) [32] to estimate the effect diverse factors from our data could have on the transformation phenotype. For the phylogenetic correlations to take into account the phylogeny, we provided the recombination-free rooted phylogenetic tree and fitted our data using the “logistic_MPLE” method with the following parameters: btol=10 and boot=100. All the test results can be found in Table S9.

### Structural prediction of hypothetical proteins of interest

To get insights into the functional role of proteins of unknown function, we characterized their protein domains. We first predicted their protein structure with *ColabFold v.1.5*. [90]. The output protein structure was then given to *Foldseek* server [91] which compared it to a large collection of protein structures (6 databases: AlphaFold/Proteome, AlphaFold/Swiss-Prot, AlphaFold/UniProt50, GMGCL, MGnify-ESM30, PDB100) in 3Di/AA mode.

### Graphical representation of GWAS output

All the graphs were done with *R v.4.1.0*.

## Acknowledgements

We thank Eloïse Guignard for technical assistance in isolation of *L. pneumophila* isolates. We thank Evelyn Skiebe for preparing and shipping *A. baumannii* isolates, and members of the RKI central sequencing lab for excellent technical assistance. We thank Eugen Pfeifer for his help in defining and providing all the data necessary for the viral quotient calculation. We are grateful to Suzana Salcedo for providing additional *A. baumannii* isolates [92]. The strains CIP and CRBIP have been obtained from the Collection Institut Pasteur (CRBIP, Paris, France). This work was supported by the French Agence Nationale de la Recherche (ANR), under grant ANR-20-CE12-0004 (TransfoConflict). EPCR lab if funded by a grant Equipe FRM (Fondation pour la Recherche Médicale) EQU201903007835 and by the Laboratoire d’Excellence IBEID Integrative Biology of Emerging Infectious Diseases [ANR-10-LABX-62-IBEID]. This work used the computational and storage services (TARS cluster) provided by the IT department at Institut Pasteur, Paris.

## References

[1] M. G. Lorenz and W. Wackernagel, “Bacterial gene transfer by natural genetic transformation in the environment.,” Microbiol. Rev., vol. 58, no. 3, Art. no. 3, Sep. 1994.

[2] O. T. Avery, C. M. MacLeod, and M. McCarty, “STUDIES ON THE CHEMICAL NATURE OF THE SUBSTANCE INDUCING TRANSFORMATION OF PNEUMOCOCCAL TYPES,” J. Exp. Med., vol. 79, no. 2, pp. 137–158, Feb. 1944.

[3] I. Chen and D. Dubnau, “DNA uptake during bacterial transformation,” Nat. Rev. Microbiol., vol. 2, no. 3, Art. no. 3, Mar. 2004, doi: 10.1038/nrmicro844.

[4] C. Johnston, B. Martin, G. Fichant, P. Polard, and J.-P. Claverys, “Bacterial transformation: distribution, shared mechanisms and divergent control,” Nat. Rev. Microbiol., vol. 12, no. 3, Art. no. 3, Mar. 2014, doi: 10.1038/nrmicro3199.

[5] D. Dubnau and M. Blokesch, “Mechanisms of DNA Uptake by Naturally Competent Bacteria,” Annu. Rev. Genet., vol. 53, no. 1, pp. 217–237, 2019, doi: 10.1146/annurev-genet-112618-043641.

[6] D. Hofreuter, S. Odenbreit, and R. Haas, “Natural transformation competence in Helicobacter pylori is mediated by the basic components of a type IV secretion system,” Mol. Microbiol., vol. 41, no. 2, pp. 379–391, 2001, doi: 10.1046/j.1365-2958.2001.02502.x.

[7] T. M. Nero, T. N. Dalia, J. C.-Y. Wang, D. T. Kysela, M. L. Bochman, and A. B. Dalia, “ComM is a hexameric helicase that promotes branch migration during natural transformation in diverse Gram-negative species,” Nucleic Acids Res., vol. 46, no. 12, pp. 6099–6111, Jul. 2018, doi: 10.1093/nar/gky343.

[8] O. H. Ambur, J. Engelstädter, P. J. Johnsen, E. L. Miller, and D. E. Rozen, “Steady at the wheel: conservative sex and the benefits of bacterial transformation,” Philos. Trans. R. Soc. B Biol. Sci., vol. 371, no. 1706, Art. no. 1706, Oct. 2016, doi: 10.1098/rstb.2015.0528.

[9] R. J. Redfield, “Genes for breakfast: the have-your-cake-and-eat-it-too of bacterial transformation,” J. Hered., vol. 84, no. 5, pp. 400–404, 1993, doi: 10.1093/oxfordjournals.jhered.a111361.

[10] B. Jeon, W. Muraoka, O. Sahin, and Q. Zhang, “Role of Cj1211 in natural transformation and transfer of antibiotic resistance determinants in Campylobacter jejuni,” Antimicrob. Agents Chemother., vol. 52, no. 8, Art. no. 8, Aug. 2008, doi: 10.1128/AAC.01607-07.

[11] K. Trzciński, C. M. Thompson, and M. Lipsitch, “Single-Step Capsular Transformation and Acquisition of Penicillin Resistance in Streptococcus pneumoniae,” J. Bacteriol., vol. 186, no. 11, pp. 3447–3452, Jun. 2004, doi: 10.1128/JB.186.11.3447-3452.2004.

[12] A.-S. Godeux et al., “Interbacterial Transfer of Carbapenem Resistance and Large Antibiotic Resistance Islands by Natural Transformation in Pathogenic Acinetobacter,” mBio, vol. 13, no. 1, pp. e02631–21, Jan. 2022, doi: 10.1128/mbio.02631-21.

[13] N. J. Croucher, R. Mostowy, C. Wymant, P. Turner, S. D. Bentley, and C. Fraser, “Horizontal DNA Transfer Mechanisms of Bacteria as Weapons of Intragenomic Conflict,” PLOS Biol., vol. 14, no. 3, Art. no. 3, Mar. 2016, doi: 10.1371/journal.pbio.1002394.

[14] A. B. Dalia, K. D. Seed, S. B. Calderwood, and A. Camilli, “A globally distributed mobile genetic element inhibits natural transformation of Vibrio cholerae,” Proc. Natl. Acad. Sci., vol. 112, no. 33, pp. 10485–10490, Aug. 2015, doi: 10.1073/pnas.1509097112.

[15] E. J. Gaasbeek et al., “A DNase Encoded by Integrated Element CJIE1 Inhibits Natural Transformation of Campylobacter jejuni,” J. Bacteriol., vol. 191, no. 7, pp. 2296–2306, Apr. 2009, doi: 10.1128/JB.01430-08.

[16] C. A. Carlson, L. S. Pierson, J. J. Rosen, and J. L. Ingraham, “Pseudomonas stutzeri and related species undergo natural transformation,” J. Bacteriol., vol. 153, no. 1, Art. no. 1, Jan. 1983, doi: 10.1128/JB.153.1.93-99.1983.

[17] B. A. Evans and D. E. Rozen, “Significant variation in transformation frequency in Streptococcus pneumoniae,” ISME J., vol. 7, no. 4, Art. no. 4, Apr. 2013, doi: 10.1038/ismej.2012.170.

[18] J. Sikorski, N. Teschner, and W. Wackernagel, “Highly different levels of natural transformation are associated with genomic subgroups within a local population of Pseudomonas stutzeri from soil,” Appl. Environ. Microbiol., vol. 68, no. 2, Art. no. 2, Feb. 2002, doi: 10.1128/aem.68.2.865-873.2002.

[19] H. Maughan and R. J. Redfield, “Extensive variation in natural competence in Haemophilus influenzae,” Evol. Int. J. Org. Evol., vol. 63, no. 7, Art. no. 7, Jul. 2009, doi: 10.1111/j.1558-5646.2009.00658.x.

[20] A.-S. Godeux et al., “Fluorescence-Based Detection of Natural Transformation in Drug-Resistant Acinetobacter baumannii,” J. Bacteriol., vol. 200, no. 19, Art. no. 19, Sep. 2018, doi: 10.1128/JB.00181-18.

[21] I. Durieux et al., “Diverse conjugative elements silence natural transformation in Legionella species,” Proc. Natl. Acad. Sci., vol. 116, no. 37, Art. no. 37, Sep. 2019, doi: 10.1073/pnas.1909374116.

[22] S. Santajit and N. Indrawattana, “Mechanisms of Antimicrobial Resistance in ESKAPE Pathogens,” BioMed Res. Int., vol. 2016, p. 2475067, 2016, doi: 10.1155/2016/2475067.

[23] A. Fodor et al., “Multidrug Resistance (MDR) and Collateral Sensitivity in Bacteria, with Special Attention to Genetic and Evolutionary Aspects and to the Perspectives of Antimicrobial Peptides—A Review,” Pathogens, vol. 9, no. 7, Art. no. 7, Jun. 2020, doi: 10.3390/pathogens9070522.

[24] V. Post and R. M. Hall, “AbaR5, a Large Multiple-Antibiotic Resistance Region Found in Acinetobacter baumannii,” Antimicrob. Agents Chemother., vol. 53, no. 6, pp. 2667–2671, Jun. 2009, doi: 10.1128/AAC.01407-08.

[25] D. W. Fraser et al., “Legionnaires’ disease: description of an epidemic of pneumonia,” N. Engl. J. Med., vol. 297, no. 22, Art. no. 22, Dec. 1977, doi: 10.1056/NEJM197712012972201.

[26] L. Gomez-Valero et al., “Extensive recombination events and horizontal gene transfer shaped the Legionella pneumophila genomes,” BMC Genomics, vol. 12, p. 536, Nov. 2011, doi: 10.1186/1471-2164-12-536.

[27] M. J. Landis and J. G. Schraiber, “Pulsed evolution shaped modern vertebrate body sizes,” Proc. Natl. Acad. Sci., vol. 114, no. 50, pp. 13224–13229, Dec. 2017, doi: 10.1073/pnas.1710920114.

[28] E. P. C. Rocha et al., “Comparisons of dN/dS are time dependent for closely related bacterial genomes,” J. Theor. Biol., vol. 239, no. 2, pp. 226–235, Mar. 2006, doi: 10.1016/j.jtbi.2005.08.037.

[29] Y. Hu, J. Zheng, and J. Zhang, “Natural Transformation in Acinetobacter baumannii W068: A Genetic Analysis Reveals the Involvements of the CRP, XcpV, XcpW, TsaP, and TonB2,” Front. Microbiol., vol. 12, p. 738034, Jan. 2022, doi: 10.3389/fmicb.2021.738034.

[30] D. Piya, L. Vara, W. K. Russell, R. Young, and J. J. Gill, “The multicomponent antirestriction system of phage P1 is linked to capsid morphogenesis,” Mol. Microbiol., vol. 105, no. 3, pp. 399–412, 2017, doi: 10.1111/mmi.13705.

[31] N. Vesel, C. Iseli, N. Guex, A. Lemopoulos, and M. Blokesch, “DNA modifications impact natural transformation of Acinetobacter baumannii,” *Nucleic Acids Res.*, p. gkad377, May 2023, doi: 10.1093/nar/gkad377.

[32] L. si Tung Ho and C. Ané, “A Linear-Time Algorithm for Gaussian and Non-Gaussian Trait Evolution Models,” Syst. Biol., vol. 63, no. 3, pp. 397–408, May 2014, doi: 10.1093/sysbio/syu005.

[33] E. P. C. Rocha and D. Bikard, “Microbial defenses against mobile genetic elements and viruses: Who defends whom from what?,” PLoS Biol., vol. 20, no. 1, p. e3001514, Jan. 2022, doi: 10.1371/journal.pbio.3001514.

[34] N. Rius, M. C. Fusté, C. Guasp, J. Lalucat, and J. G. Lorén, “Clonal Population Structure of Pseudomonas stutzeri, a Species with Exceptional Genetic Diversity,” J. Bacteriol., vol. 183, no. 2, pp. 736–744, Jan. 2001, doi: 10.1128/JB.183.2.736-744.2001.

[35] M. Vos, “Why do bacteria engage in homologous recombination?,” Trends Microbiol., vol. 17, no. 6, pp. 226–232, Jun. 2009, doi: 10.1016/j.tim.2009.03.001.

[36] L. Hadany and T. Beker, “On the evolutionary advantage of fitness-associated recombination.,” Genetics, vol. 165, no. 4, pp. 2167–2179, Dec. 2003.

[37] J. Chen et al., “Genome hypermobility by lateral transduction,” Science, vol. 362, no. 6411, pp. 207–212, Oct. 2018, doi: 10.1126/science.aat5867.

[38] P. H. Oliveira, M. Touchon, J. Cury, and E. P. C. Rocha, “The chromosomal organization of horizontal gene transfer in bacteria,” Nat. Commun., vol. 8, no. 1, Art. no. 1, Oct. 2017, doi: 10.1038/s41467-017-00808-w.

[39] M. L. Gwinn, R. Ramanathan, H. O. Smith, and J.-F. Tomb, “A New Transformation-Deficient Mutant of Haemophilus influenzae Rd with Normal DNA Uptake,” J. Bacteriol., vol. 180, no. 3, pp. 746–748, Feb. 1998.

[40] P. Jorth and M. Whiteley, “An Evolutionary Link between Natural Transformation and CRISPR Adaptive Immunity,” mBio, vol. 3, no. 5, pp. e00309–12, Oct. 2012, doi: 10.1128/mBio.00309-12.

[41] R. J. Redfield, W. A. Findlay, J. Bossé, J. S. Kroll, A. D. Cameron, and J. H. Nash, “Evolution of competence and DNA uptake specificity in the Pasteurellaceae,” BMC Evol. Biol., vol. 6, no. 1, p. 82, Oct. 2006, doi: 10.1186/1471-2148-6-82.

[42] S. Stutzmann and M. Blokesch, “Comparison of chitin-induced natural transformation in pandemic Vibrio cholerae O1 El Tor strains,” Environ. Microbiol., vol. 22, no. 10, pp. 4149–4166, 2020, doi: 10.1111/1462-2920.15214.

[43] G. Wilharm, J. Piesker, M. Laue, and E. Skiebe, “DNA Uptake by the Nosocomial Pathogen Acinetobacter baumannii Occurs during Movement along Wet Surfaces,” J. Bacteriol., vol. 195, no. 18, pp. 4146–4153, Sep. 2013, doi: 10.1128/JB.00754-13.

[44] A.-S. Godeux et al., “Scarless Removal of Large Resistance Island AbaR Results in Antibiotic Susceptibility and Increased Natural Transformability in Acinetobacter baumannii,” Antimicrob. Agents Chemother., vol. 64, no. 10, Sep. 2020, doi: 10.1128/AAC.00951-20.

[45] M. Haudiquet, J. M. de Sousa, M. Touchon, and E. P. C. Rocha, “Selfish, promiscuous and sometimes useful: how mobile genetic elements drive horizontal gene transfer in microbial populations,” Philos. Trans. R. Soc. B Biol. Sci., vol. 377, no. 1861, p. 20210234, Aug. 2022, doi: 10.1098/rstb.2021.0234.

[46] L. Attaiech et al., “Silencing of natural transformation by an RNA chaperone and a multitarget small RNA,” Proc. Natl. Acad. Sci. U. S. A., vol. 113, no. 31, pp. 8813–8818, Aug. 2016, doi: 10.1073/pnas.1601626113.

[47] F. Tesson, A. Hervé, M. Touchon, C. d’Humières, J. Cury, and A. Bernheim, “Systematic and quantitative view of the antiviral arsenal of prokaryotes,” Sep. 2021. doi: 10.1101/2021.09.02.458658.

[48] C. Berndt, P. Meier, and W. Wackernagel, “DNA restriction is a barrier to natural transformation in Pseudomonas stutzeri JM300,” Microbiology, vol. 149, no. 4, pp. 895–901, 2003, doi: 10.1099/mic.0.26033-0.

[49] O. Humbert, M. S. Dorer, and N. R. Salama, “Characterization of Helicobacter pylori factors that control transformation frequency and integration length during inter-strain DNA recombination,” Mol. Microbiol., vol. 79, no. 2, pp. 387–401, 2011, doi: 10.1111/j.1365-2958.2010.07456.x.

[50] J. P. Holt, A. J. Grant, C. Coward, D. J. Maskell, and J. J. Quinlan, “Identification of Cj1051c as a Major determinant for the restriction barrier of Campylobacter jejuni StrainNCTC11168,” Appl. Environ. Microbiol., vol. 78, no. 22, pp. 7841–7848, 2012, doi: 10.1128/AEM.01799-12.

[51] J. M. Beauchamp, R. M. Leveque, S. Dawid, and V. J. DiRita, “Methylation-dependent DNA discrimination in natural transformation of Campylobacter jejuni,” Proc. Natl. Acad. Sci. U. S. A., vol. 114, no. 38, pp. E8053–E8061, 2017, doi: 10.1073/pnas.1703331114.

[52] M. B. Finn, K. M. Ramsey, H. J. Tolliver, S. L. Dove, and M. R. Wessels, “Improved transformation efficiency of group A Streptococcus by inactivation of a type I restriction modification system,” PloS One, vol. 16, no. 4, p. e0248201, 2021, doi: 10.1371/journal.pone.0248201.

[53] K. A. Jolley and M. C. Maiden, “BIGSdb: Scalable analysis of bacterial genome variation at the population level,” BMC Bioinformatics, vol. 11, no. 1, p. 595, Dec. 2010, doi: 10.1186/1471-2105-11-595.

[54] K. L. Wyres, S. M. Cahill, K. E. Holt, R. M. Hall, and J. J. Kenyon, “Identification of Acinetobacter baumannii loci for capsular polysaccharide (KL) and lipooligosaccharide outer core (OCL) synthesis in genome assemblies using curated reference databases compatible with Kaptive,” *Microb*. Genomics, vol. 6, no. 3, p. e000339, 2020, doi: 10.1099/mgen.0.000339.

[55] S. M. Cahill, R. M. Hall, and J. J. Kenyon, “An update to the database for Acinetobacter baumannii capsular polysaccharide locus typing extends the extensive and diverse repertoire of genes found at and outside the K locus,” *Microb*. Genomics, vol. 8, no. 10, p. 000878, 2022, doi: 10.1099/mgen.0.000878.

[56] S. K. Gupta et al., “ARG-ANNOT, a new bioinformatic tool to discover antibiotic resistance genes in bacterial genomes,” Antimicrob. Agents Chemother., vol. 58, no. 1, pp. 212–220, 2014, doi: 10.1128/AAC.01310-13.

[57] B. Jia et al., “CARD 2017: expansion and model-centric curation of the comprehensive antibiotic resistance database,” Nucleic Acids Res., vol. 45, no. D1, pp. D566–D573, Jan. 2017, doi: 10.1093/nar/gkw1004.

[58] E. Doster et al., “MEGARes 2.0: a database for classification of antimicrobial drug, biocide and metal resistance determinants in metagenomic sequence data,” Nucleic Acids Res., vol. 48, no. D1, pp. D561–D569, Jan. 2020, doi: 10.1093/nar/gkz1010.

[59] L. Chen, D. Zheng, B. Liu, J. Yang, and Q. Jin, “VFDB 2016: hierarchical and refined dataset for big data analysis--10 years on,” Nucleic Acids Res., vol. 44, no. D1, pp. D694–697, Jan. 2016, doi: 10.1093/nar/gkv1239.

[60] A. Perrin and E. P. C. Rocha, “PanACoTA: a modular tool for massive microbial comparative genomics,” NAR Genomics Bioinforma., vol. 3, no. lqaa106, Art. no. lqaa106, Feb. 2021, doi: 10.1093/nargab/lqaa106.

[61] C. Camacho et al., “BLAST+: architecture and applications,” BMC Bioinformatics, vol. 10, p. 421, 2009, doi: 10.1186/1471-2105-10-421.

[62] L. A. Ronish, E. Lillehoj, J. K. Fields, E. J. Sundberg, and K. H. Piepenbrink, “The structure of PilA from Acinetobacter baumannii AB5075 suggests a mechanism for functional specialization in Acinetobacter type IV pili,” J. Biol. Chem., vol. 294, no. 1, pp. 218–230, Jan. 2019, doi: 10.1074/jbc.RA118.005814.

[63] P. Cingolani et al., “A program for annotating and predicting the effects of single nucleotide polymorphisms, SnpEff,” Fly (Austin), vol. 6, no. 2, pp. 80–92, Apr. 2012, doi: 10.4161/fly.19695.

[64] J. T. L. Kang and N. A. Rosenberg, “Mathematical properties of linkage disequilibrium statistics defined by normalization of the coefficient D=pAB−pApB,” Hum. Hered., vol. 84, no. 3, pp. 127– 143, 2019, doi: 10.1159/000504171.

[65] K. Katoh and D. M. Standley, “MAFFT Multiple Sequence Alignment Software Version 7: Improvements in Performance and Usability,” Mol. Biol. Evol., vol. 30, no. 4, pp. 772–780, Apr. 2013, doi: 10.1093/molbev/mst010.

[66] L.-T. Nguyen, H. A. Schmidt, A. von Haeseler, and B. Q. Minh, “IQ-TREE: A Fast and Effective Stochastic Algorithm for Estimating Maximum-Likelihood Phylogenies,” Mol. Biol. Evol., vol. 32, no. 1, pp. 268–274, Jan. 2015, doi: 10.1093/molbev/msu300.

[67] S. Kalyaanamoorthy, B. Q. Minh, T. K. F. Wong, A. von Haeseler, and L. S. Jermiin, “ModelFinder: fast model selection for accurate phylogenetic estimates,” Nat. Methods, vol. 14, no. 6, Art. no. 6, Jun. 2017, doi: 10.1038/nmeth.4285.

[68] D. T. Hoang, O. Chernomor, A. von Haeseler, B. Q. Minh, and L. S. Vinh, “UFBoot2: Improving the Ultrafast Bootstrap Approximation,” Mol. Biol. Evol., vol. 35, no. 2, pp. 518–522, Feb. 2018, doi: 10.1093/molbev/msx281.

[69] N. J. Croucher et al., “Rapid phylogenetic analysis of large samples of recombinant bacterial whole genome sequences using Gubbins,” Nucleic Acids Res., vol. 43, no. 3, Art. no. 3, Feb. 2015, doi: 10.1093/nar/gku1196.

[70] L. J. Revell, “phytools: an R package for phylogenetic comparative biology (and other things),” Methods Ecol. Evol., vol. 3, no. 2, pp. 217–223, 2012, doi: 10.1111/j.2041-210X.2011.00169.x.

[71] S. A. Fritz and A. Purvis, “Selectivity in Mammalian Extinction Risk and Threat Types: a New Measure of Phylogenetic Signal Strength in Binary Traits,” Conserv. Biol., vol. 24, no. 4, pp. 1042– 1051, 2010.

[72] M. W. Pennell et al., “geiger v2.0: an expanded suite of methods for fitting macroevolutionary models to phylogenetic trees,” Bioinformatics, vol. 30, no. 15, pp. 2216–2218, Aug. 2014, doi: 10.1093/bioinformatics/btu181.

[73] S. A. Ishikawa, A. Zhukova, W. Iwasaki, and O. Gascuel, “A Fast Likelihood Method to Reconstruct and Visualize Ancestral Scenarios,” Mol. Biol. Evol., vol. 36, no. 9, Art. no. 9, Sep. 2019, doi: 10.1093/molbev/msz131.

[74] S. S. Abby, B. Néron, H. Ménager, M. Touchon, and E. P. C. Rocha, “MacSyFinder: A Program to Mine Genomes for Molecular Systems with an Application to CRISPR-Cas Systems,” PLOS ONE, vol. 9, no. 10, Art. no. 10, Oct. 2014, doi: 10.1371/journal.pone.0110726.

[75] B. Neron, R. Denise, C. Coluzzi, M. Touchon, E. P. C. Rocha, and S. S. Abby, “MacSyFinder v2: Improved modelling and search engine to identify molecular systems in genomes.” bioRxiv, p. 2022.09.02.506364, Feb. 28, 2023. doi: 10.1101/2022.09.02.506364.

[76] J. Cury, M. Touchon, and E. P. C. Rocha, “Integrative and conjugative elements and their hosts: composition, distribution and organization,” Nucleic Acids Res., vol. 45, no. 15, pp. 8943–8956, Sep. 2017, doi: 10.1093/nar/gkx607.

[77] Z. Xie and H. Tang, “ISEScan: automated identification of insertion sequence elements in prokaryotic genomes,” Bioinformatics, vol. 33, no. 21, Art. no. 21, Nov. 2017, doi: 10.1093/bioinformatics/btx433.

[78] J. Guo et al., “VirSorter2: a multi-classifier, expert-guided approach to detect diverse DNA and RNA viruses,” Microbiome, vol. 9, no. 1, Art. no. 1, Feb. 2021, doi: 10.1186/s40168-020-00990-y.

[79] S. Nayfach, A. P. Camargo, F. Schulz, E. Eloe-Fadrosh, S. Roux, and N. C. Kyrpides, “CheckV assesses the quality and completeness of metagenome-assembled viral genomes,” Nat. Biotechnol., vol. 39, no. 5, Art. no. 5, May 2021, doi: 10.1038/s41587-020-00774-7.

[80] P. Terzian et al., “PHROG: families of prokaryotic virus proteins clustered using remote homology,” NAR Genomics Bioinforma., vol. 3, no. 3, p. lqab067, Sep. 2021, doi: 10.1093/nargab/lqab067.

[81] B. Néron, E. Littner, M. Haudiquet, A. Perrin, J. Cury, and E. P. C. Rocha, “IntegronFinder 2.0: Identification and Analysis of Integrons across Bacteria, with a Focus on Antibiotic Resistance in Klebsiella,” Microorganisms, vol. 10, no. 4, Art. no. 4, Apr. 2022, doi: 10.3390/microorganisms10040700.

[82] E. P. Nawrocki and S. R. Eddy, “Infernal 1.1: 100-fold faster RNA homology searches,” Bioinforma. Oxf. Engl., vol. 29, no. 22, pp. 2933–2935, Nov. 2013, doi: 10.1093/bioinformatics/btt509.

[83] S. R. Eddy, “Accelerated Profile HMM Searches,” PLOS Comput. Biol., vol. 7, no. 10, p. e1002195, Oct. 2011, doi: 10.1371/journal.pcbi.1002195.

[84] D. Hyatt, G.-L. Chen, P. F. LoCascio, M. L. Land, F. W. Larimer, and L. J. Hauser, “Prodigal: prokaryotic gene recognition and translation initiation site identification,” BMC Bioinformatics, vol. 11, no. 1, p. 119, Mar. 2010, doi: 10.1186/1471-2105-11-119.

[85] D. Couvin et al., “CRISPRCasFinder, an update of CRISRFinder, includes a portable version, enhanced performance and integrates search for Cas proteins,” Nucleic Acids Res., vol. 46, no. W1, pp. W246–W251, Jul. 2018, doi: 10.1093/nar/gky425.

[86] I. Grissa, G. Vergnaud, and C. Pourcel, “CRISPRFinder: a web tool to identify clustered regularly interspaced short palindromic repeats,” Nucleic Acids Res., vol. 35, no. suppl_2, pp. W52–W57, Jul. 2007, doi: 10.1093/nar/gkm360.

[87] A. Biswas, J. N. Gagnon, S. J. J. Brouns, P. C. Fineran, and C. M. Brown, “CRISPRTarget: bioinformatic prediction and analysis of crRNA targets,” RNA Biol., vol. 10, no. 5, pp. 817–827, May 2013, doi: 10.4161/rna.24046.

[88] J. A. Lees, M. Galardini, S. D. Bentley, J. N. Weiser, and J. Corander, “pyseer: a comprehensive tool for microbial pangenome-wide association studies,” Bioinformatics, vol. 34, no. 24, Art. no. 24, Dec. 2018, doi: 10.1093/bioinformatics/bty539.

[89] Y. Benjamini and Y. Hochberg, “Controlling the False Discovery Rate: A Practical and Powerful Approach to Multiple Testing,” J. R. Stat. Soc. Ser. B Methodol., vol. 57, no. 1, pp. 289–300, 1995, doi: 10.1111/j.2517-6161.1995.tb02031.x.

[90] M. Mirdita, K. Schütze, Y. Moriwaki, L. Heo, S. Ovchinnikov, and M. Steinegger, “ColabFold: making protein folding accessible to all,” Nat. Methods, vol. 19, no. 6, Art. no. 6, Jun. 2022, doi: 10.1038/s41592-022-01488-1.

[91] M. van Kempen, et al., “Foldseek: fast and accurate protein structure search.” bioRxiv, p. 2022.02.07.479398, Sep. 20, 2022. doi: 10.1101/2022.02.07.479398.

[92] T. Rubio et al., “Incidence of an Intracellular Multiplication Niche among Acinetobacter baumannii Clinical Isolates,” mSystems, vol. 7, no. 1, pp. e00488–21, doi: 10.1128/msystems.00488-21.

