## Supplemental Figures for "Intragenomic conflicts with plasmids and chromosomal mobile genetic elements drive the evolution of natural transformation within species"

Fanny Mazzamurro[1,2,+], Jason Baby Chirakadavil[3], Isabelle Durieux[3], Ludovic Poiré[3], Julie Plantade[3], Christophe Ginevra[4], Sophie Jarraud[4], Gottfried Wilharm[5], Xavier Charpentier[3,*], Eduardo P. C. Rocha[1,+*]
[1] Institut Pasteur, Université Paris Cité, CNRS UMR3525, Microbial Evolutionary Genomics, Paris, 75015, France.
[2] Collège Doctoral – Sorbonne Université, F-75005 Paris, France – France
[3] CIRI, Centre International de Recherche en Infectiologie – Inserm, U1111, Université Claude Bernard Lyon 1, CNRS, UMR5308, École Normale Supérieure de Lyon, Univ Lyon, 69100, Villeurbanne, France – France
[4] Centre national de Référence des Légionelles – Centre de biologie Nord, 103 grande-rue de la Croix-Rousse 69317 LYON Cedex 04, France – France
[5] Robert Koch Institute, Project group P2, Wernigerode Branch, D-38855 Wernigerode, Germany

+ co-corresponding authors
* co-senior authors

**Table S1** List of phylogenetic trees computed in this study and their features

**Table S2** Measures of phylogenetic signal on continuous transformation rates

**Table S3** Evolutionary models tested on the log10-transformed transformation rates and their fitness in *Legionella pneumophila* and *Acinetobacter baumannii*

**Table S4** Association of transformation phenotype and change with recombination features tested with Wilcoxon tests

**Table S5** a. GWAS results obtained with a unitig approach on the binary transformation phenotype in *Legionella pneumophila*
b. GWAS results obtained with a unitig approach on the binary transformation phenotype in *Acinetobacter baumannii*

**Table S6** Collection of *Legionella pneumophila* and *Acinetobacter baumannii* strains and their measures of transformation rates

**Table S7** Reproducibility of the transformation luminescence assay in *Legionella pneumophila* and *Acinetobacter baumannii*

**Table S8** a. Protein-encoding genes involved in natural transformation in *Legionella pneumophila*
b. sncRNA genes involved in natural transformation in *Legionella pneumophila*
c. Protein-encoding genes involved in natural transformation in *Acinetobacter baumannii*

**Table S9** Statistical tests performed throughout the study comparing transformable and non-transformable strains in *Legionella pneumophila* and *Acinetobacter baumannii*

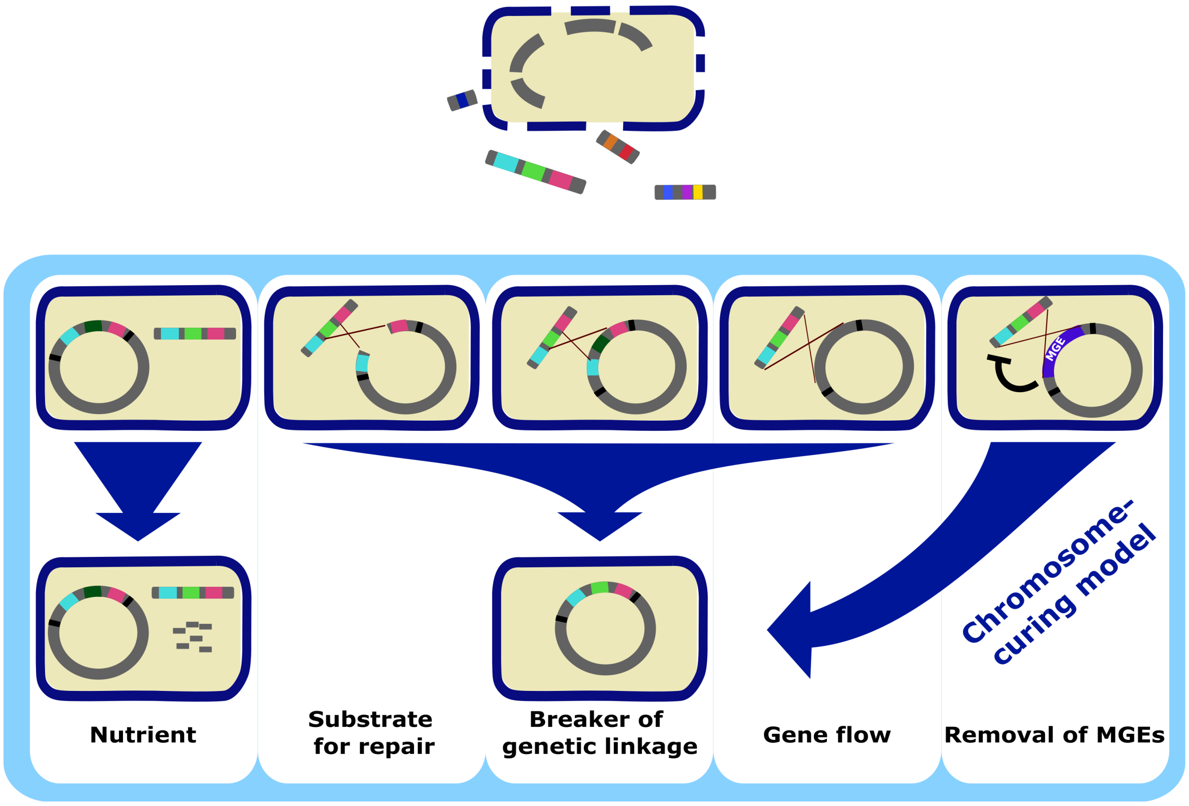


**Figure S1 Evolutionary theories behind natural transformation's persistence**

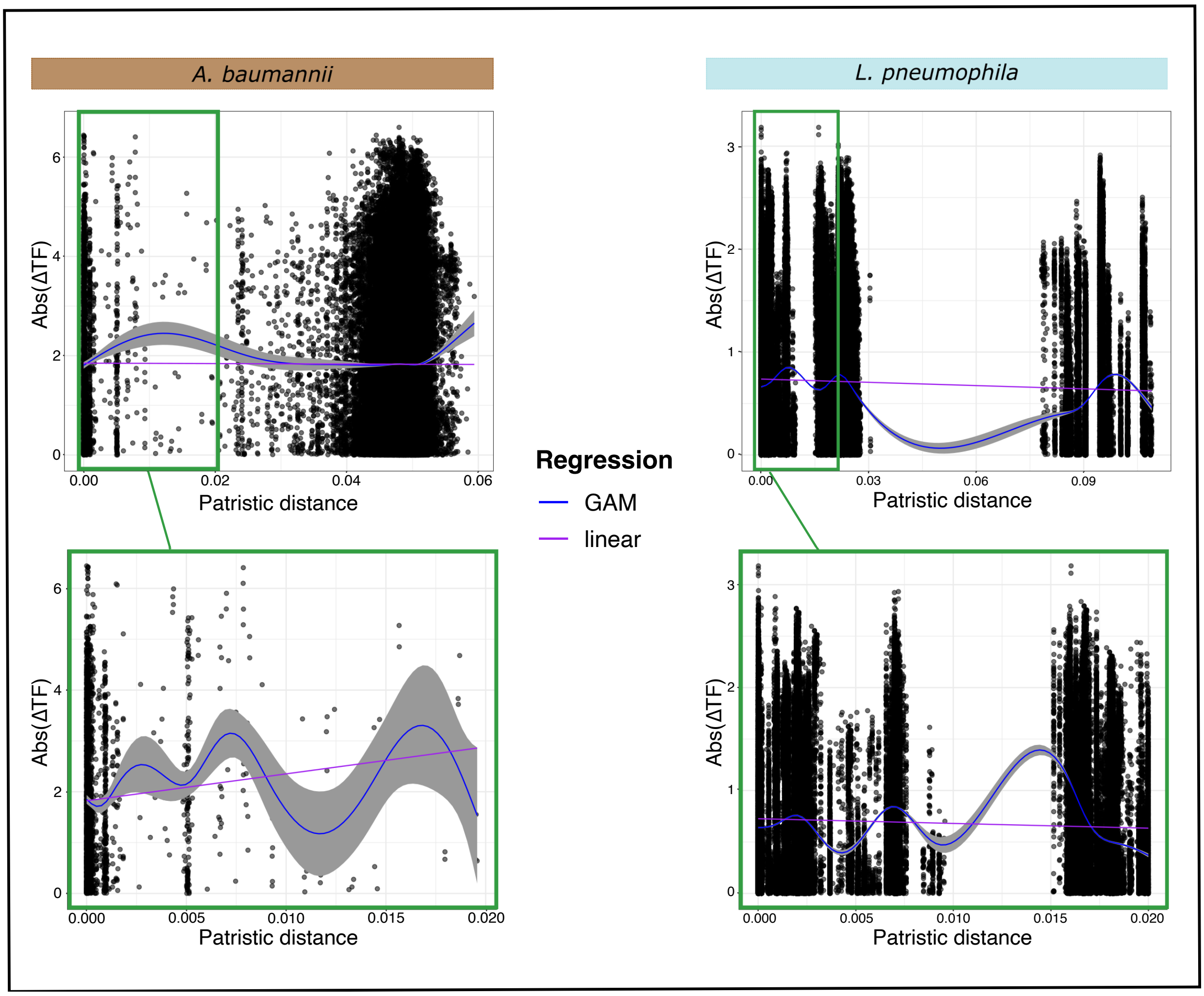


**Figure S2 Absolute variation of transformation rates between two strains depending on their patristic distances in *Acinetobacter baumanni* (left) and *Legionella pneumophila* (right).** Regressions between abs(ΔTF) and patristic distance were performed with a linear model (purple line) and with a generalized additive model (GAM default parameters; blue line).

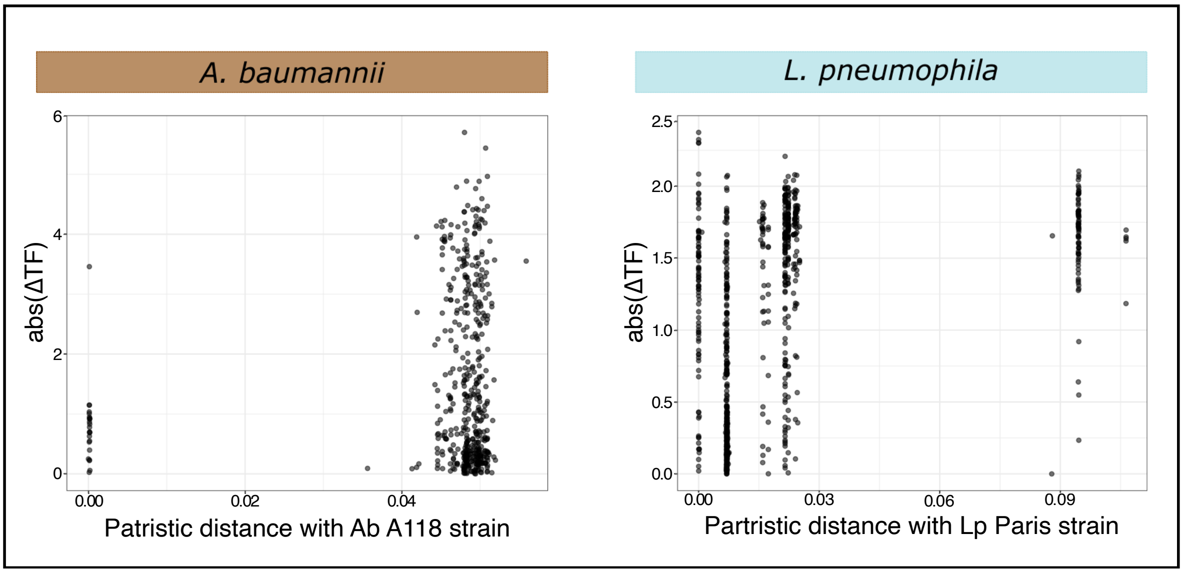


**Figure S3 Variation of strain transformation rate depending on its patristic distance with the donor strain in *Acinetobacter baumannii* (left) and *Legionella pneumophila* (right)**

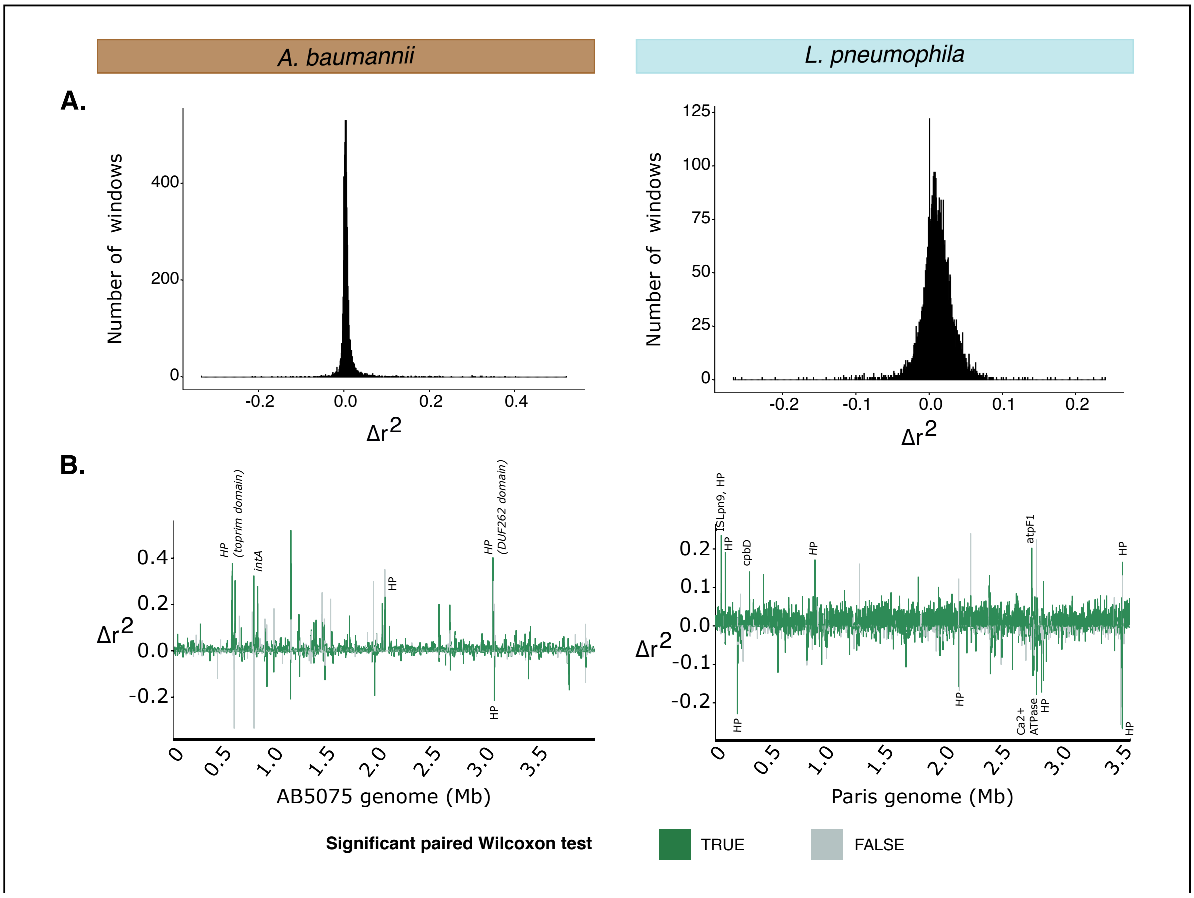


**Figure S4 Difference between transformable and non-transformable strains of their squared correlation ($r^2$) between bi-allelic values at two loci in windows of 500 nt along their genome**

A. Distribution of $\Delta r^2$ in the 500 nt screened windows in *Acinetobacter baumannii* (left) and *Legionella pneumophila* (right). $\Delta r^2$ was calculated as $r^2_{mean}(NT)-r^2_{mean}(T)$

b. Distribution of $\Delta r^2$ along the reference genome divided into 500 nt windows in *Acinetobacter baumannii* (left) and *Legionella pneumophila* (right): AB5075 for Ab, Paris for Lp. The windows in which the distribution of $r^2$ between transformable and non-transformable populations was significantly different according to a paired Wilcoxon test were colored in green, otherwise they were grey. The highest peaks of $\Delta r^2$ were annotated with the genes the corresponding window was overlapping. HP stands for hypothetical proteins.

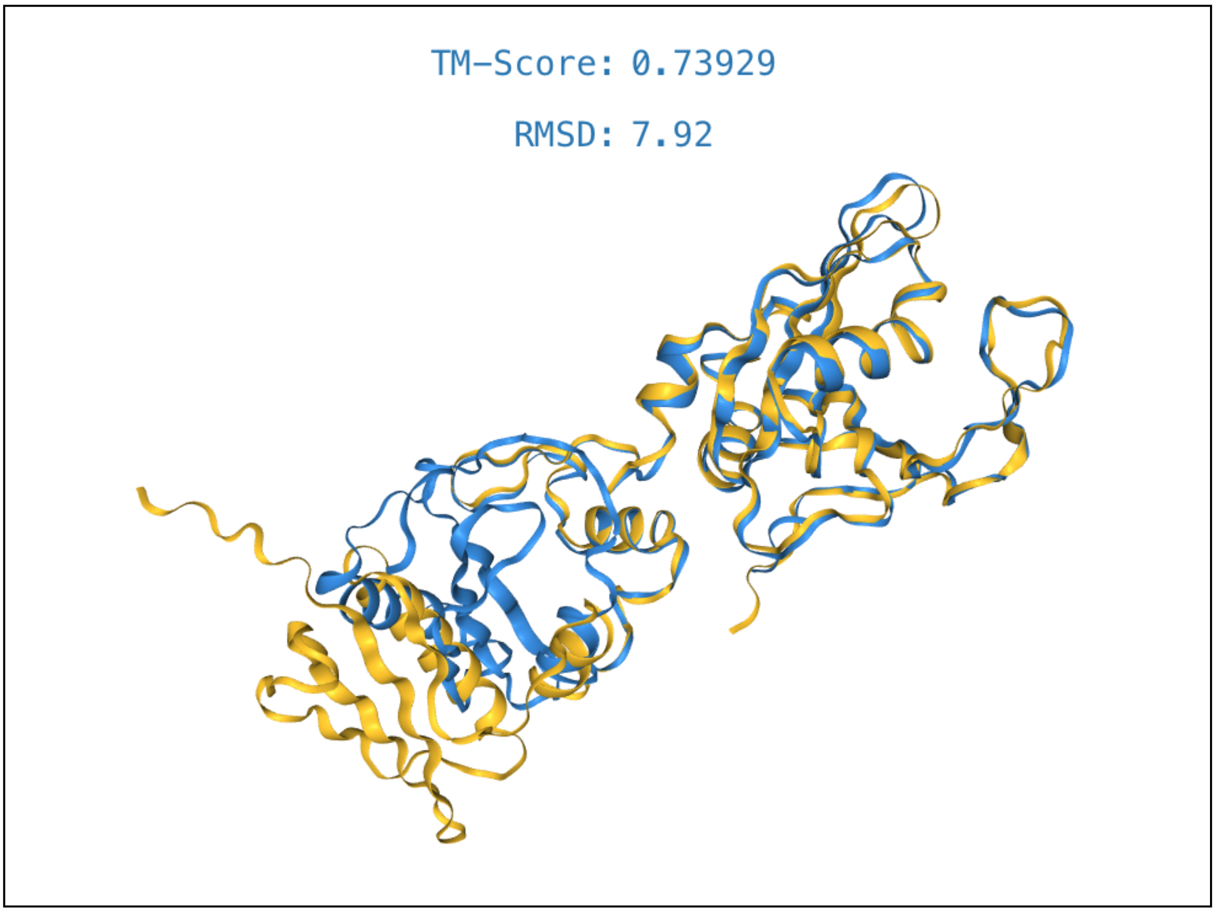


**Figure S5 Alignment of the structure of the DarA_N domain-containing protein in *A. baumannii* ACICU (A0A4Y3J949 UniProt) (yellow) with the structure of the most significantly associated protein (pangenome family 6781) with non-transformability carried by prophages (blue) in *Acinetobacter baumannii***

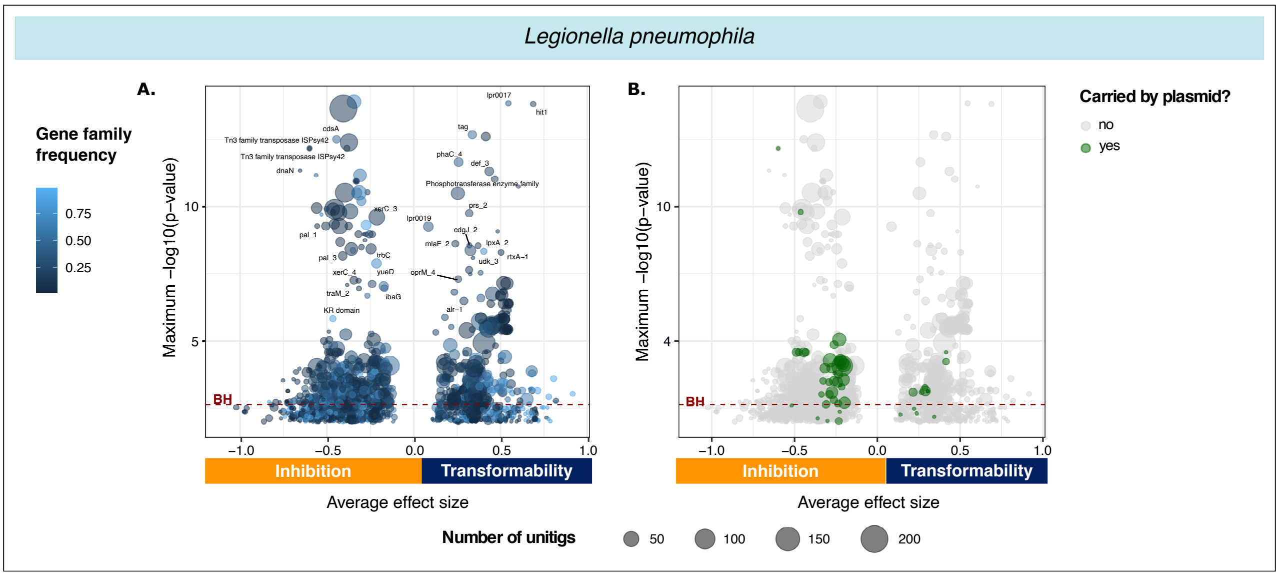


**Figure S6 Volcano plots showing average effect sizes and significance of the association of the gene families with the transformation phenotype according to $GWAS_U^{bin}$-cov in *Legionella pneumophila* and plasmid association with the phenotype.** Each circle stands for a gene family. The size of the circle depends on the number of unitigs that mapped the gene in all the samples. The value on the x-axis corresponds to the average effect size of all the unitigs mapping the gene. The y-axis indicates how significant this effect can be by representing the maximal -log10-transformed p-value adjusted for population structure of all the unitigs of this gene. Significantly associated gene families are above the Benjamini-Hochberg (BH) threshold (red dashed line).

A. Gene families were colored by their frequencies in the collection.

B. Gene families that were carried by a plasmid were colored in green.

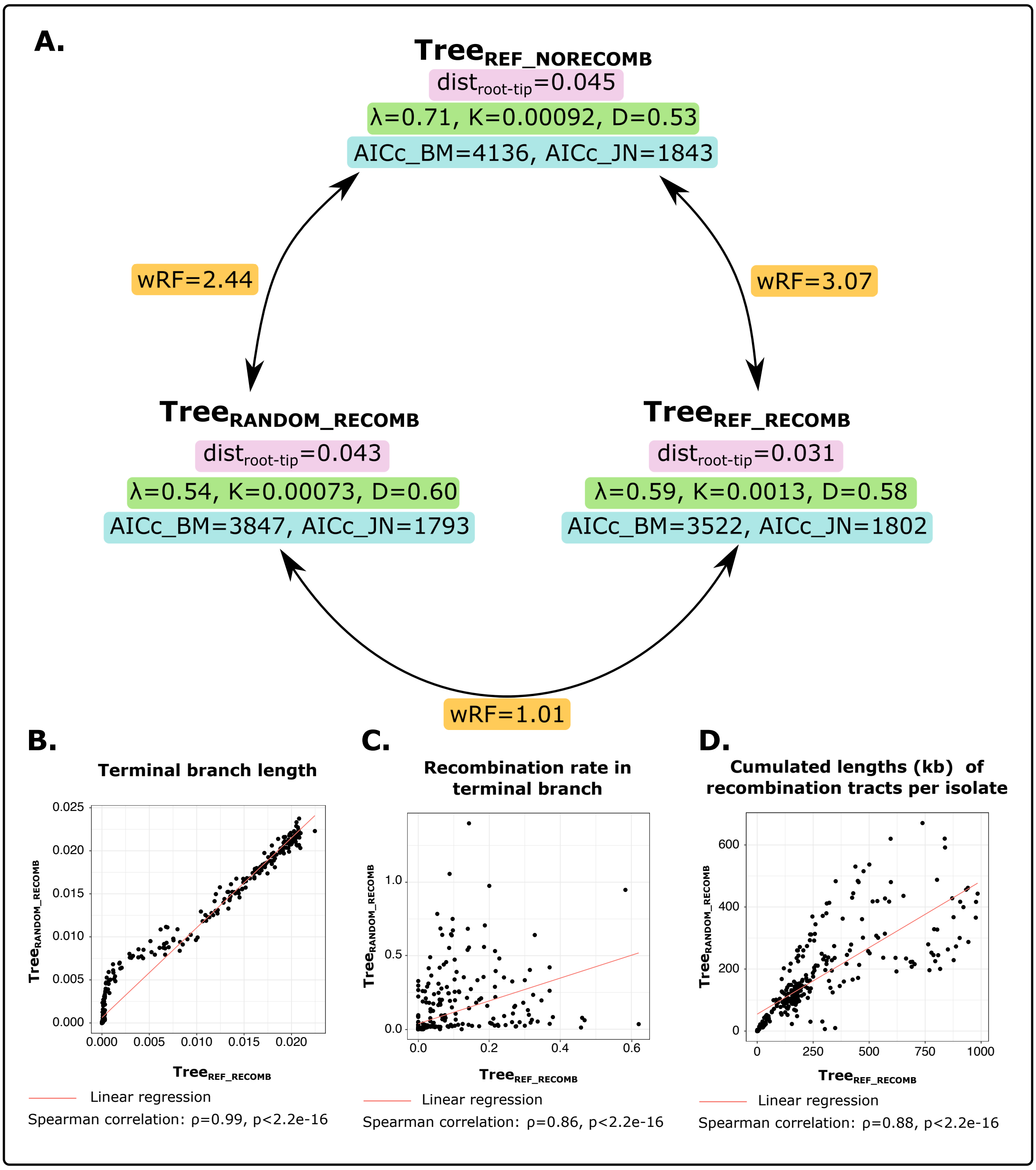


**Figure S7 Comparison of the different phylogenies inferred in *Acinetobacter baumannii* based on different evolutionary features.** $\text{Tree}_{\text{REF\_NORECOMB}}$ is the tree inferred without taking into account recombination on the alignment of concatenated persistent genes ordered according to a complete reference genome (AB5075 genome). $\text{Tree}_{\text{REF\_RECOMB}}$ is the tree inferred taking into account recombination on the alignment of concatenated persistent genes ordered according to a complete reference genome (AB5075 genome). $\text{Tree}_{\text{RANDOM\_NORECOMB}}$ is the tree inferred on the alignment of concatenated persistent genes randomly ordered and that takes into account recombination.

A. Comparison of different evolutionary features in regards to one another and to the transformation phenotype: topological distance (weighted Robinson-Foulds distance wRF), average distance root to tip ($\text{dist}_{\text{root-tip}}$), phylogenetic signal of the transformation phenotype, fit of the trait to the Brownian motion (BM) and Jump Node (JN) evolutionary models.

B. Pairwise comparison of the terminal branch lengths when provided with an ordered alignment of concatenated persistent genes or a random one.

C. Pairwise comparison of the recombination rate *Gubbins* estimated when provided with an ordered alignment of concatenated persistent genes or a random one.

D. Pairwise comparison of the cumulated lengths of recombination tracts identified by *Gubbins* when provided with an ordered alignment of concatenated persistent genes or a random one.

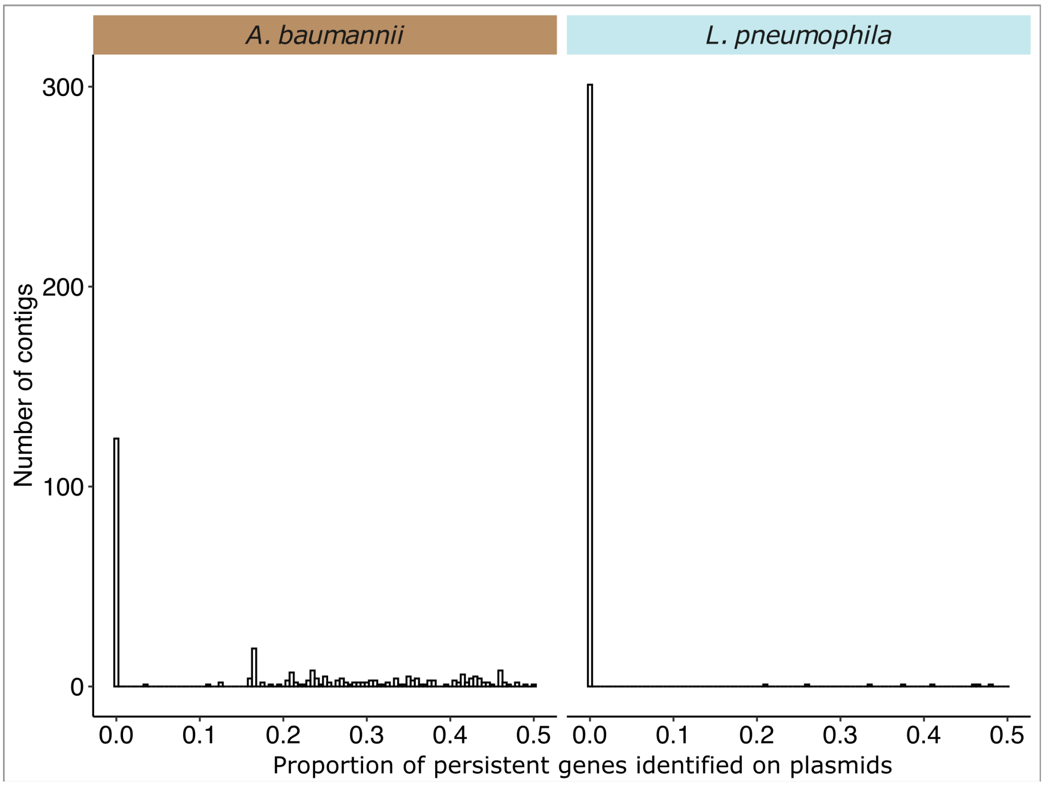


**Figure S8 Distribution of the proportion of persistent genes in contigs assigned as plasmids in *Acinetobacter baumannii* (left) and *Legionella pneumophila* (right)**
